# The State of Motion: Demographic and Representational Consequences of Motion-Based Exclusion in FMRI

**DOI:** 10.64898/2026.07.23.739855

**Authors:** Patrick Sadil, Briha Ansari, Agostina Casamento-Moran, Ann S. Choe, Jungin Choi, Farzad V. Farahani, Lukman E. Ismaila, Micah Alan Johnson, Arunkumar Kannan, Mary Beth Nebel, James J. Pekar, Haris Iqbal Sair, Joshua Stim, Adrian M. Svingos, Tor D. Wager, Martin A. Lindquist

**Affiliations:** Johns Hopkins Bloomberg School of Public Health; Department of Biomedical Engineering, Johns Hopkins School of Medicine; Kennedy Krieger Institute; Department of Radiology, The Russell H. Morgan Department of Radiology and Radiological Science, Johns Hopkins University School of Medicine; Kirby Research Center for Functional Brain Imaging, Kennedy Krieger Institute; Johnson & Johnson; Department of Electrical and Computer Engineering, Johns Hopkins Whiting School of Engineering; Department of Neurology, Johns Hopkins School of Medicine; Center for Neurodevelopmental and Imaging Research, Kennedy Krieger Institute; The Malone Center for Engineering in Healthcare, The Whiting School of Engineering, Johns Hopkins University; Brain Injury Clinical Research Center, Kennedy Krieger Institute; Department of Physical Medicine and Rehabilitation, Johns Hopkins University School of Medicine; Department of Psychological and Brain Sciences, Dartmouth College

**Keywords:** MRI

## Abstract

Large-scale neuroimaging datasets are increasingly used to map relationships between brain structure, function, and behavior across the human lifespan. Routinely, analyses exclude participants who moved too much during imaging. While this decision is framed as quality control, it is increasingly recognized that head motion is not randomly distributed across individuals within a study, and so motion-based exclusion may preferentially remove people with particular characteristics relevant to the scientific goals of the study. Here we survey head motion and how it relates to participant characteristics across six large, publicly available datasets spanning nearly the entire human lifespan, namely the Human Connectome Project (HCP) Young Adult, HCP in Development, HCP in Aging, Adolescent Brain Cognitive Development^SM^ Study, UK Biobank, and Spatial Topology project. These six datasets comprise more than 50,000 unique participants and 300,000 scans. We further benchmark our findings against motion distributions aggregated by MRIQC across more than 1.5 million scans. Under commonly applied strict exclusion thresholds, large fractions of participants would be removed (exceeding 80 % in the UK Biobank task data), and these removals were demographically structured, disproportionately excluding younger and older participants, those with higher BMI, and those with motion-associated clinical conditions. Respiratory pseudo-motion inflated estimates of head motion in adult cohorts, and applying notch filtering to remove respiratory frequencies from these estimates meaningfully reduced exclusion rates. Exclusion also carried downstream consequences. Strict thresholds reduced statistical power, inflated study costs, and altered the apparent predictability of behavioral phenotypes by removing a non-random, behaviorally distinct subgroup. These findings demonstrate that motion exclusion thresholds are not neutral quality-control decisions but structured selection mechanisms that reshape the composition of neuroimaging samples. We recommend that studies report the demographic characteristics of excluded participants, prefer data-driven censoring methods over fixed motion cutoffs, and clarify the target population while considering appropriate weighting techniques.

## 1 Introduction

Functional magnetic resonance imaging (fMRI) studies routinely exclude data from participants who move when scanned. This decision is framed as quality control, based on the well-established finding that head motion introduces artifacts into fMRI data through multiple mechanisms, including signal intermixing and spin history effects (Friston et al., 1996; Maknojia et al., 2019). These artifacts can take multiple forms and interact in complex ways with both acquisition parameters and participant neuroanatomy (Faraji-Dana et al., 2016b; Faraji-Dana et al., 2016a; Havsteen et al., 2017), and failure to account for them can substantially bias results (Chyzhyk et al., 2022; Makowski et al., 2019; Nebel et al., 2022; Power et al., 2012; Siegel et al., 2017; Van Dijk et al., 2012; Weinberger & Radulescu, 2016). To mitigate these effects, numerous prospective and retrospective methods have been developed (Chyzhyk et al., 2022; Ciric et al., 2017; Maknojia et al., 2019; Muschelli et al., 2014; Parkes et al., 2018; Pruim et al., 2015; Ramduny et al., 2025; Reddy et al., 2023; White et al., 2010; Yan et al., 2013; Yang et al., 2024). However, no existing approach fully eliminates motion-related bias (Power et al., 2020), and exclusion of high-motion participants remains common. Motion is thus not only a source of image degradation but also a potential source of systematic sample selection, a consequence that has received far less attention. The scale of participant exclusion is considerable. For example, in two reports using data from the Adolescent Brain Cognitive Development study, 40 % and 60 % of participants were excluded (Marek et al., 2019, 2022). Such high rates of exclusion are perhaps unsurprising in a pediatric cohort, especially one enriched with children exhibiting externalizing symptoms that may be associated with difficulty holding still (Barch et al., 2018; Cosgrove et al., 2022; Garavan et al., 2018; Siegel et al., 2017). However, high exclusion rates are not limited to developmental samples. The Human Connectome Project Young Adult dataset is widely regarded as a high-quality resource (Marcus et al., 2013), yet analyses that apply a strict criterion to mitigate motion effects on functional connectivity have excluded more than half of the participants (Greene et al., 2018). These examples suggest that commonly used motion thresholds can remove large fractions of participants, even in carefully curated datasets.

Unfortunately, participant motion correlates systematically with demographic and clinical characteristics (Cosgrove et al., 2022; Makowski et al., 2019), meaning that motion-based exclusion functions as a non-random filter on study samples (Peverill et al., 2025). For example, children with more profound clinical symptoms of autism spectrum disorder often move more in the scanner than their neurotypical peers or even children with less profound symptoms, potentially yielding analytic samples that underrepresent this population (Nebel et al., 2022; Shi et al., 2025). Elevated motion relative to healthy controls has also been reported in individuals with multiple sclerosis, schizophrenia, bipolar disorder, and stroke survivors (Pardoe et al., 2016; Seto et al., 2001; Wylie et al., 2014; Yao et al., 2017). Motion levels are stable within individuals across scans (Kong et al., 2014; Liu et al., 2017; Van Dijk et al., 2012; Yan et al., 2013; Zeng et al., 2014; Zuo et al., 2014), and they has been associated with a broad range of characteristics in the general population, including age, sex, body size, hyperactivity-impulsivity, blood pressure, language ability, and nicotine use, among other factors (Bolton et al., 2020; Ekhtiari et al., 2019; Frew et al., 2022; Kong et al., 2014; Power et al., 2020; Satterthwaite et al., 2013; Shi et al., 2025; Siegel et al., 2017; Teo et al., 2024; Ward et al., 2024).

Consider the case of the ABCD Study®, which, to our knowledge, is the only population-level dataset that has been used to explore these relationships between motion, participant characteristics, motion-based exclusion, and biased downstream inferences (Peverill et al., 2025). One of the strengths of the ABCD study design is the use of multi-stage probability sampling, which resulted in a sample population that approximately represents the US (Garavan et al., 2018). Although this approximate matching is imperfect (Compton et al., 2019; Gard et al., 2023) and also conflicts with other goals of the ABCD, such as the recruitment of a relatively high proportion of children who exhibit symptoms related to mental health issues, a sample population that closely resembles the target (US) population along major axes like race and sex simplifies some analyses. However, excluding participants based on motion alters the distribution of even those characteristics that resemble the target population, like race and ethnicity (Cosgrove et al., 2022). For example, non-white children were approximately twice as likely as white children to be excluded based on motion. In some circumstances, such losses can be mitigated by statistical techniques (Gard et al., 2023; LeWinn et al., 2017; Nebel et al., 2022). But there remains a need to understand the prevalence of these issues in datasets beyond the ABCD, to understand how routine preprocessing decisions shape representativeness and, ultimately, to understand the generalizability of conclusions drawn from population neuroimaging studies.

Here, we sought to understand the scope of these issues across the neuroimaging field, exploring the extent to which motion acts as a population-level filter (that is, a non-random selection mechanism that determines which participants remain available for analyses). To do this, we survey head motion across six prominent, publicly available datasets spanning childhood through older adulthood: the Human Connectome Project-Young Adult (HCPYA Marcus et al., 2013), UK Biobank (UKB Alfaro-Almagro et al., 2018), Human Connectome Project in Development (HCPD Harms et al., 2018), Human Connectome Project in Aging (HCPA Harms et al., 2018), the Adolescent Brain Cognitive Development (ABCD Casey et al., 2018) study, and the Spatial Topology (SpaceTop) project (Jung et al., 2025). Together, these datasets comprise more than 50,000 unique participants and more than 300,000 scans. We further benchmark our findings against motion distributions reported by MRIQC (Esteban et al., 2017), which, as of writing, aggregates data from over 1.5 million scans. By quantifying motion across large-scale cohorts, we assess both its magnitude and its demographic consequences under commonly applied exclusion thresholds. We address four questions (Table 1): (A) How much do participants move across major datasets? (B) In these datasets, which participant characteristics are related to motion? (C) Which demographic groups are disproportionately excluded? and (D) What are the implications for sample composition and downstream inference? We find that exclusion is both substantial and demographically patterned, with negative implications for the generalizability of population neuroimaging findings.

**Table 1.** Results Overview. Analyses aim to answer four main questions. Several of the findings reported here have been reported elsewhere (e.g., relationships with motion and BMI); here, we show that they are present in several large-scale datasets (ABCD, HCPYA, HCPD, HCPA, UKB, SpaceTop), and that in these datasets there remains an unresolved tension between biasing study results by including low-quality (that is, motion-corrupted) data and biasing study results by excluding high-motion participants.

| Topic | Result |
| --- | --- |
| How much motion? | • Motion varies across datasets, including those with overlapping demographics (Figure 1, Figure 9).<br>• Motion follows consistent temporal patterns (Figure 2 through Figure 7).<br>– In resting-state fMRI, motion increases throughout scanning.<br>– In task fMRI, motion correlates with task timing. |
| Who moves? | • Within participants, motion is consistent across scans and sessions, even those that span years (Figure 8).<br>• The oldest and youngest participants tend to move more, as do those with higher BMI (Figure 9).<br>• Motion co-varies with several psychological and Life Function variables (Figure 10). |
| Who is excluded? | • When using common exclusion criteria, rates of exclusion can be substantial (e.g., >80 %) (Figure 11, Figure 12).<br>• Excluding participants based on motion substantially alters the sex, age, and BMI composition of datasets (Figure 13). |
| Do the exclusions matter? | • Even with modern denoising techniques, motion substantially changes functional connectivity (Figure 14).<br>• Motion-related exclusion increases study costs and decreases model predictive performance (Figure 15).<br>• Statistical power in Brain-wide Association Studies is minimally affected by motion-based exclusion for the largest datasets, but reduced by 25 % in smaller (600–800 participant) datasets that sample from high-motion demographics (HCPA, HCPD, Figure 16).<br>• Predictive models trained with high-motion participants achieve higher levels of performance than those trained with low-motion participants (Figure 17). |

## 2 Methods

### 2.1 Datasets

We extracted sociodemographic, clinical, and fMRI data from six publicly available datasets. Across these datasets, several types of tasks were employed. As part of this survey, we examined the temporal properties of motion, hypothesizing that they would vary with task design. Below, we briefly summarize the design of each study. For details of each task, please see the cited references.

From each dataset, participants were excluded from the analyses if key demographic variables were unavailable. For demographics, we focused on three that were generally available: age, sex, and body mass index (BMI). When a variable was unavailable across the entire dataset, we note this explicitly below. Additional dataset-specific exclusion criteria are listed below. Note that these exclusions differ from the motion-based exclusions that are the primary subject of this survey.

#### 2.1.1 The HCP Young Adult (HCPYA) 1200 Subjects Data Release

The HCPYA comprises 1097 participants aged 22–37 years. Whole-brain EPI data were acquired on a Siemens Skyra 3 T scanner at Washington University in St. Louis with TR=720 ms, TE=33.1 ms, flip angle=52^◦^, 2 mm isotropic voxels, 72 slices, and a multi-band acceleration factor of 8. For each task, two runs were acquired with opposite phase-encoding directions (LR and RL). Full acquisition details are provided in (Feinberg et al., 2010; Moeller et al., 2010; Setsompop et al., 2012; Xu et al., 2012).

Seven tasks were administered, each designed to evoke well-characterized activity in specific neural systems (Barch et al., 2013): an emotion face-matching task, a card-guessing reward task, a story/arithmetic language task, a motor task involving finger tapping, toe squeezing, and tongue movement, a relational reasoning task, a social cognition task using animated shapes, and a working memory N-back task. For each task, two runs were acquired; displays of motion by time retained task-block structure when timing was consistent across participants. Full task details are provided in (Barch et al., 2013). Participants also completed four resting-state fMRI runs, each approximately 14.5 min in duration, acquired with alternating phase-encoding directions (LR and RL).

The HCPYA dataset provides the variable gender for each participant, which we refer to throughout this report as sex.

**Exclusions.** Participants were excluded for being flagged for exhibiting any of several of the quality control issues identified by the Human Connectome Project (154 participants with either anatomical anomalies, head coil instabilities, or rfMRI artifact).

#### 2.1.2 The HCP in Development (HCPD)

The HCPD Release 2.0 includes 652 participants aged 5–21 years, with imaging data collected at four sites (Harvard, University of California, Los Angeles, Washington University in St. Louis, and the University of Minnesota). All MRI data were acquired using 3 T Siemens Prisma scanners (Siemens, Erlangen, Germany). Participants between 5- and 7-years old were scanned using a pediatric 32-channel head coil developed by Ceresensa (ceresensa.com); participants between 8- and 21-years old were scanned using the Siemens 32-channel Prisma head coil. Full acquisition details are provided in (Harms et al., 2018).

Three tasks were administered (Somerville et al., 2018): a reward guessing task with consistent block timing (Guessing), a go/no-go inhibitory control task with jittered timing (CARIT), and a face/shape emotion-matching task adapted from the HCPYA (Emotion). For 5- to 7-year-old participants, six resting-state fMRI runs of approximately 3.5 min each were acquired (alternating AP/PA phase encoding); for 8- to 21-year-old participants, four runs of approximately 6.5 min each were acquired. For several analyses, we split the HCPD by age cohort (5–7 vs. 8+).

**Exclusions.** 12 participants were excluded due to anatomical brain anomalies identified by a radiologist (as provided by HCP) or missing BMI data.

#### 2.1.3 Human Connectome Project-Aging (HCPA)

The HCPA dataset includes 725 participants aged 36–100. Ages in the HCPA dataset were right-censored to protect participants’ privacy; all ages above 89 are reported as 100. Three tasks were administered (Bookheimer et al., 2019): a go/no-go inhibitory control task identical to the HCPD version (CARIT), a paired-associates face-name memory task with encoding and recall blocks (FaceName), and a visuomotor task involving button presses in response to flickering checkerboard stimuli (VisMotor). Resting-state fMRI comprised four runs of approximately 6.5 min each, acquired with alternating phase-encoding directions. Full task details are provided in (Harms et al., 2018).

**Exclusions.** 27 participants were excluded due to anatomical brain anomalies identified by a radiologist (as provided by HCP) or missing BMI information.

#### 2.1.4 Adolescent Brain Cognitive Development (ABCD)

The ABCD study is an ongoing longitudinal study of more than 10,000 9- to 10-year-old children in the United States through adolescence, with neuroimaging data collected every two years. The dataset was enriched with children who exhibited internalizing and externalizing symptoms. In the release accessed here (7.0), there were 11458 participants. For a detailed description of recruitment, demographics, and study exclusion rules, see (Barch et al., 2018; Garavan et al., 2018; Thompson et al., 2019). For a detailed description of the acquisition, see the report by (Casey et al., 2018). Our analyses focused on data curated as a part of the ABCD Community Collection (Feczko et al., 2021).

Three tasks were administered: a monetary incentive delay task with jittered trial timing (MID Knutson et al., 2000), a stop-signal task in which response timing is determined by performance (SST Logan, 1994), and an emotional N-back task with consistent block timing (EN-back Barch et al., 2013). Resting-state fMRI comprised four runs of 5 min each; participants were instructed to fixate on a crosshair with eyes open.

As a part of our analyses of the ABCD dataset, we categorized ABCD participants as exhibiting externalizing, internalizing, or attention problems. These categories were based on the normed scores for the Brief Problem Monitor (Achenbach, 2009), using a cutoff of ≥ 65 (averaged across sessions for which the score was available).

Unique to the ABCD dataset, the motion parameters were not filtered for this report (see Section 2.4). Instead, we used the filtered motion traces as provided in the ABCD Community Collection.

**Exclusions.** 96 participants were excluded due to missing demographics from at least one session, or when all runs were flagged by the official ABCD quality control procedures (Hagler Jr et al., 2019). Specifically, we excluded scans that did not satisfy the ABCD inclusion criteria, excluding those criteria that were related to censored frames (inclusion: fMRI raw QC, T1 raw QC, behavior passed, E-prime timing match OR ignore E-prime mismatch, fMRI B0 unwarp available, FreeSurfer QC not failed, fMRI manual post-processing QC not failed, fMRI registration to T1w, fMRI dorsal cutoff score, fMRI ventral cutoff score, derived results exist). Additionally, all resting-state scans with run above 4 or task scans with run above 2 were excluded, and all runs were truncated to the modal length by task. This resulted in the exclusion of 5908 out of 206071 scans.

#### 2.1.5 UK Biobank (UKB)

The UKB imaging subsample includes participants from a large prospective epidemiological cohort (Alfaro-Almagro et al., 2018). In the accessed release, there were 44659 participants with usable imaging data. A single task was administered—a modified version of the HCPYA face/shape emotion-matching task (Hariri et al., 2002), with identical timing across all participants—alongside a single resting-state scan of approximately 6 min.

**Exclusions.** We excluded 1188 participants for missing BMI data.

#### 2.1.6 Spatial Topology (SpaceTop)

The SpaceTop dataset includes 117 young adult participants with six hours of fMRI data combined across six tasks and four sessions, all with jittered timing (Jung et al., 2025). The tasks were designed to span a broad range of cognitive and affective processes: an emotionally salient video-viewing task with continuous affective ratings (align-video), a multimodal social cognition task involving somatic pain, vicarious pain, and mental rotation (social), a narrative comprehension task with ratings of affect and expectation (narratives), a dynamic face perception and rating task (faces), a self-referential theory-of-mind task based on previously viewed video characters (shortvideo), and a fractionated task comprising attention reorienting, memory encoding/retrieval, and two theory-of-mind paradigms (fractional). Full descriptions of each task, including trial structure and timing, are provided in (Jung et al., 2025).

**Exclusions.** BMI data were unavailable for this dataset, and so BMI was not used to exclude participants. All participants had complete data on age and sex.

### 2.2 Quantifying Motion using Framewise Displacement

To quantify motion, we used framewise displacement (FD), which measures the displacement of a point on the cortical surface between two successive time points (Power et al., 2012). FD is computed from the six motion parameters obtained during realignment (three translational and three rotational), with the rotational parameters converted to millimeters by calculating the arc-length displacement on a sphere of radius 50 mm. The displacement at time *t* is then the sum of the absolute temporal first differences of the six motion parameters:

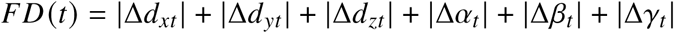

where Δ denotes the difference operator (i.e., Δ*β*_*t*_ = *β*_*t*_ − *β*_*t*−1_). The variables *d*_*xt*_, *d*_*yt*_, and *d*_*zt*_ denote translations, and *α*_*t*_, *β*_*t*_, and *γ*_*t*_ denote rotations, each measured relative to a reference volume. By convention, the FD at time *t* = 0 is set equal to zero and excluded from all analyses.

### 2.3 Defining Framewise Displacement Thresholds for Unacceptable Motion

Motion-based exclusion criteria vary with the analysis, the effects of interest, and the overall data quality. Common approaches include thresholding on mean FD, requiring a minimum number of frames to be below a target displacement, or flagging scans with large, isolated motion spikes. The appropriate values for these three criteria are vociferously debated but generally depend on the analysis and specific features of the dataset in question.

For our exclusion analyses, we applied two regimes, as described by (Kay et al., 2025; Parkes et al., 2018). The “lenient” regime excluded participants whose average FD exceeded 0.55 mm. The “strict” regime applied three criteria simultaneously, excluding participants who met any of the following conditions: mean FD exceeding 0.25 mm, more than 20 % of frames with FD above 0.2 mm, or maximum FD exceeding 5 mm.

### 2.4 Filtered Motion

Common thresholds for framewise displacement (e.g., 0.15 mm to 0.2 mm, 0.5 mm) were developed before sub-second TRs became common (Power et al., 2012; Power et al., 2014). At higher sampling rates made feasible by multislice acquisition sequences, elevated framewise displacement shows less correspondence with data-driven measures of signal atypicality, such as carpet plots, DVARS, or projection scrubbing (Fair et al., 2020; Phạm et al., 2023; Power et al., 2019). One possible explanation is that respiration alters the magnetic field in a way that causes the head to appear to move—a phenomenon termed “pseudo-motion”—without substantially impacting data quality through spin history effects (Fair et al., 2020; Van De Moortele et al., 2002). Fair et al. (2020) therefore argued that motion parameters used to assess data quality should be filtered to remove frequencies associated with respiration (around 0.3 Hz by adolescence).

Across all analyzed datasets, motion traces exhibited clear power at frequencies associated with respiration (Section A.2). Respiratory frequencies had the highest power in translation parameters aligned with the phase-encoding direction, consistent with the hypothesis that this apparent motion is driven in part by magnetic-field changes rather than by purely mechanical movement. We therefore repeated analyses after removing respiration-related frequencies from the motion parameters.

In each scan condition, respiration frequencies were estimated from the translation parameter corresponding to the phase-encoding axis. Motion parameters were linearly detrended, and the spectrum was estimated with a multitaper filter (number of points: 512; half bandwidth: 8). The respiration frequency was identified as the peak frequency in the range 0.1 Hz to 0.6 Hz. Peak frequencies were averaged across scans within participants for each scan condition. To remove respiratory-related frequencies, we adopted an approach similar to the “Fixed Frequency” method of (Fair et al., 2020). The lower and upper cutoff frequencies of a second-order infinite-impulse-response notch filter were set at the median and third quartile of the distribution of peak frequencies within each condition and session. Although peak frequencies were estimated from the motion parameters rather than from direct respiratory recordings, the resulting notch filter closely resembled the respiration band filter reported by Fair et al. (2020) (Table 1).

**Table 1.**
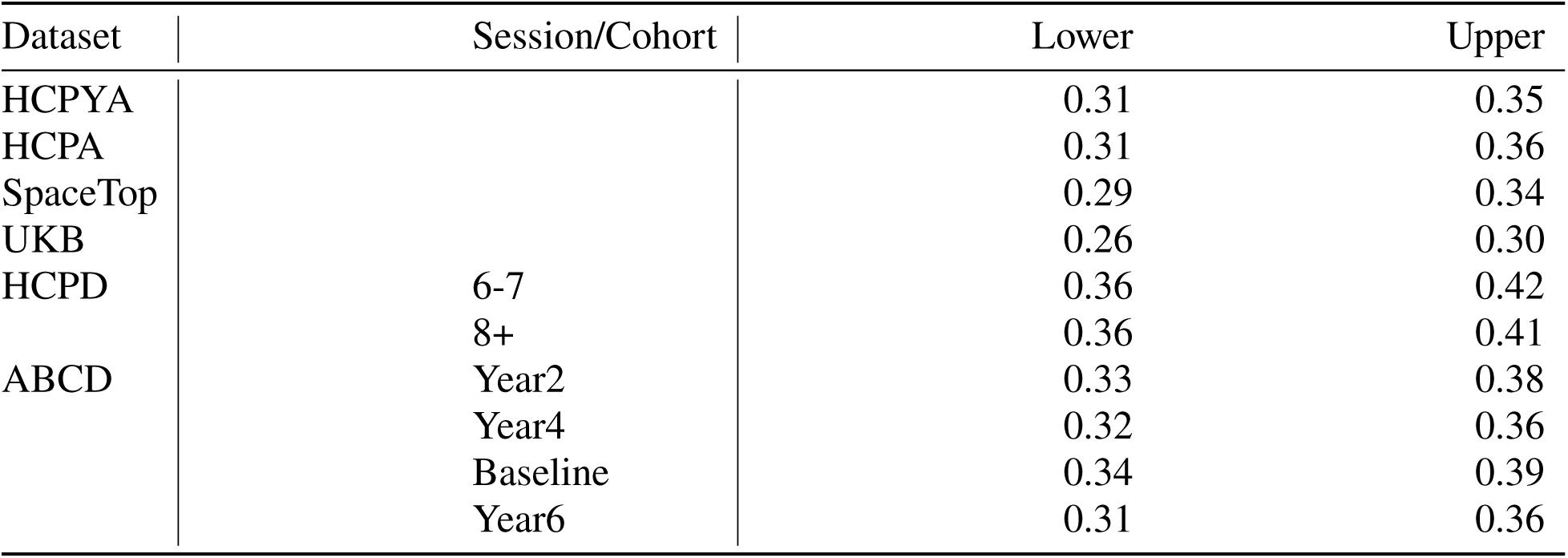
Summary of Parameters Used for Notch Filtering. Each entry indicates the average (across conditions) upper and lower frequency for the filter designed to remove respiration frequencies from motion parameters. These limits are comparable to the range observed in a subset of ABCD participants (N=62) using a respiratory band: 0.31 Hz to 0.43 Hz (Fair et al., 2020). Note that the motion parameters for the ABCD were filtered as part of the ABCD Community Collection (Feczko et al., 2021).

| Dataset | Session/Cohort | Lower | Upper |
| --- | --- | --- | --- |
| HCPYA |  | 0.31 | 0.35 |
| HCPA |  | 0.31 | 0.36 |
| SpaceTop |  | 0.29 | 0.34 |
| UKB |  | 0.26 | 0.30 |
| HCPD | 6-7 | 0.36 | 0.42 |
|  | 8+ | 0.36 | 0.41 |
| ABCD | Year2 | 0.33 | 0.38 |
|  | Year4 | 0.32 | 0.36 |
|  | Baseline | 0.34 | 0.39 |
|  | Year6 | 0.31 | 0.36 |

### 2.5 Classifying High- and Low-Movers

To examine factors related to motion and assess the stability of motion as a trait, we classified participants as “high-movers” or “low-movers.” Classification was performed using a simple median split of the mean framewise displacement from a single index scan for the datasets in which this analysis was conducted. The index scan for HCPYA was REST1 LR (Session 1), and the first resting-state run of the baseline session for ABCD.

### 2.6 Assessing the Consistency of Head Motion Within and Across Sessions

To assess the reliability of motion within sessions, we measured the intraclass correlation coefficient (ICC) for average FD in the resting-state scans of the HCP datasets using the `irr` package (Gamer et al., 2019). We used a single unit, two-way random-effects measure of consistency: ICC(C,1).

To assess the reliability of motion across sessions separated by years, we measured the ICC for average FD in the resting-state scans of the ABCD and UKB datasets.

### 2.7 Characterizing the Impact of Motion-Related Data Removal on Downstream Analyses

To determine how data exclusion affects results, we conducted two complementary analyses. First, we sought to determine whether motion-based exclusions are justified with modern preprocessing techniques by replicating findings showing that poor data quality (e.g., inclusion of motion-corrupted frames) biases results (Section 2.7.1). Second, we sought to demonstrate that, even though inclusion of these participants introduces bias, excluding them also negatively affects a study’s results (Section 2.9).

#### 2.7.1 Evaluating Changes in Functional Connectivity on Preprocessed Data

To estimate the extent to which including motion-riddled data can negatively impact results, we measured the mean absolute change (MAC) in functional connectivity (Williams et al., 2022). The MAC is a quantitative extension of a qualitative, graphical method presented by (Power et al., 2014), who assessed the impact of motion on connectivity across inter-nodal distances (see also Fair et al., 2020). The MAC measures the average change in connectivity estimates when censoring is based on data quality, compared with random censoring. Frames are first ordered by a quality metric (e.g., framewise displacement). A fixed proportion of the worst frames is censored, and functional connectivity estimates are calculated with the censored timeseries.

Surrogate timeseries are then generated by randomly shuffling the data quality labels. Those timeseries are censored and used to calculate surrogate connectivity estimates. For each pair of nodes, the distribution of absolute differences between the original connectivity estimate and the surrogate connectivity estimate is calculated and summarized with an average. The process is then repeated across exclusion thresholds and participants.

This measure is based on the idea that poor data quality biases connectivity estimates. If removing a fixed proportion of a scan based on data quality results in a greater change in functional connectivity than removing the same proportion at random, then quality-based censoring yields less-biased estimates. We used this measure to show that even scans that have undergone extensive preprocessing remain substantially biased by the inclusion of high-motion (that is, low-quality) data.

To calculate the MAC, we used a 500-participant subset of the UKB. Following (Williams et al., 2022), 10 surrogate timeseries were calculated at each level of thresholding. For these analyses, we considered three preprocessing strategies: (1) the standard UKB preprocessing stream (fix+smooth, see Alfaro-Almagro et al., 2018), (2) regression-based denoising comprising 24 motion parameters (Friston et al., 1996), 4 spatial parameters (CSF, WM, and their first-order differences), and bandpass filtering (Butterworth order 5, 0.08 Hz to 0.9 Hz), and (3) bandpass filtering only. In all cases, datasets were spatially smoothed using a 6 mm FWHM Gaussian kernel. Connectivities were estimated using the 7 Network Schaefer 100 atlas (Mansour et al., 2023; Schaefer et al., 2018). Data quality was assessed using framewise displacement and DPD measured in the uncleaned (immediately before denoising) scans (defined below in Section 2.8).

### 2.8 DVARS

With modern scanning parameters, several reports suggest that motion measures are not accurate indicators of data quality, and that data-driven methods are preferred (Fair et al., 2020; Phạm et al., 2023; Power et al., 2019). One common data-driven measure is DVARS, or the spatial root-mean-square of the change in the signal. This is used to assess temporal changes in signal variance across brain voxels between consecutive brain volumes. High DVARS values indicate substantial atypicality in the signal and are often associated with head motion, physiological noise, or other artifacts. Frames are removed when they exceed pre-specified thresholds. Many thresholds are used in practice, depending on how DVARS is normalized. Here we rely on formulations and recommendations provided by Afyouni and Nichols (2018).

In the following, each scan consists of *T* time points (brain volumes/frames), each with *V* voxels. Let *x*_*vt*_ denote the fMRI signal at voxel *v* and time *t*, *m*_*v*_ the temporal mean at voxel *v*, and *m* the grand mean across all voxels and time points. To begin, we center and scale the data as follows:

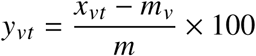

The total (all) variation at time *t* is defined as A-var:

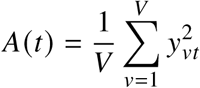

and the variance of the first difference as D-var:

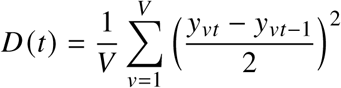

DVARS is defined as:

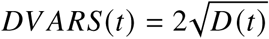

It is often of interest to determine whether this value is significantly different from some value *μ*_0_. Assuming normality, the corresponding *χ*^2^-statistic is of the form:

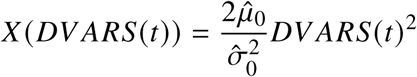

Under the null *X* (*DV ARS*(*t*)) follows an approximate 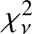 with 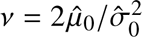 degrees of freedom. Here *μ*^_0_ is the mean of 2*D*(*t*) under the null, and 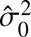 is the corresponding variance. For methods of estimating *μ̂*_0_ and 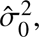 see Afyouni and Nichols (2018). The values of *X* (*DV ARS*(*t*)) can be converted to *p*-values, denoted *P*(*DV ARS*(*t*)), and then to *z*-scores:

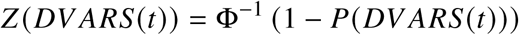

where Φ^−1^(·) represents the inverse of the cumulative distribution function of the standard normal. *Z* (*DV ARS*(*t*)) corresponds to statistical significance. Practical significance can be measured with the change in percent D-var from baseline:

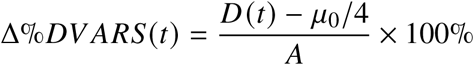

where *A* is the average value of *A*(*t*). We refer to this quantity as DPD and use it as our primary censoring metric.

### 2.9 Costs of Including High-Motion Participants on Study Budget and Model Performance

We quantified the expected impact of motion on design budgets and predictive performance using the economic framework presented by Ooi et al. (2025). The framework is based on a model that estimates prediction performance, assessed as the correlation between true and predicted values, *ρ*, from a study’s sample size, *n*, and per-participant scan time, 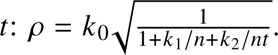 The *k*_*i*_ parameters relate to both maximum performance and the relative rates at which increasing sample size or scan time approaches that maximum. Ooi et al. estimated the *k*_*i*_ from existing datasets. By augmenting this model with information about per-minute scan costs and per-participant recruitment costs, the framework links study design directly to budgetary constraints. To achieve a target usable sample size *N*, the required recruitment sample is estimated as 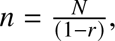 where *r* is the exclusion rate.

To evaluate sampling bias within the Ooi et al. framework, we assessed differences in prediction performance across low- and high-motion HCPYA participants. We selected HCP participants for this analysis because of their uniquely long runs and constructed two partially overlapping groups of 800 participants each, defined by the lowest and highest average framewise displacement.

Using these groups, we trained models to predict several behavioral phenotypes from connectivity matrices, subsetting the available data to vary both the number of participants and the scan length used to compute connectivity. We focus on the 18 phenotypes examined by Ooi et al. to follow the model underlying their framework.

### 2.10 Statistical Reporting Conventions

For reporting parameter estimates, we follow the recommendations of Louis and Zeger (2008) and report the estimate with 95 % confidence intervals in surrounding subscripts.

## 3 Results

### 3.1 How Much Do Participants Move?

Across all studies, participants moved an average of approximately 0.2 mm between frames, with motion varying substantially across datasets and scan types. Motion tended to be highest in the ABCD dataset (Figure 1), particularly in the baseline session where participants were approximately 9-10 years old. Average FD in the ABCD baseline session often exceeded 0.21 mm. In contrast, the UKB and HCPYA datasets showed much lower average motion, typically below 0.15 mm. This aligns with well-established findings that children and adolescents exhibit substantially more head motion than adults. In subsequent sessions (Year 2 and Year 4), differences between the ABCD cohort and the adult datasets (HCPYA, UKB) were reduced. Average head motion in the SpaceTop dataset occupies an intermediate range, with notable variability across its diverse task battery. Together, these findings indicate that motion magnitude varies substantially across large-scale studies and depends strongly on both age and study design. In general, motion tended to increase throughout a session, such that later runs showed more motion than earlier ones (Figure 2 through Figure 7). For example, in all three ABCD sessions, average motion increased progressively across resting-state runs. Likewise, in the first session of SpaceTop, each subsequent social run exhibited higher average motion than the previous one. This progressive increase suggests fatigue, reduced compliance, or discomfort as plausible contributors to within-session motion drift. However, this pattern did not hold for the resting-state scans in the HCPD and HCPA datasets, where the second resting-state run often had somewhat less motion than the first, suggesting that session structure and scheduling may modulate temporal patterns of movement.

**Figure 1.**
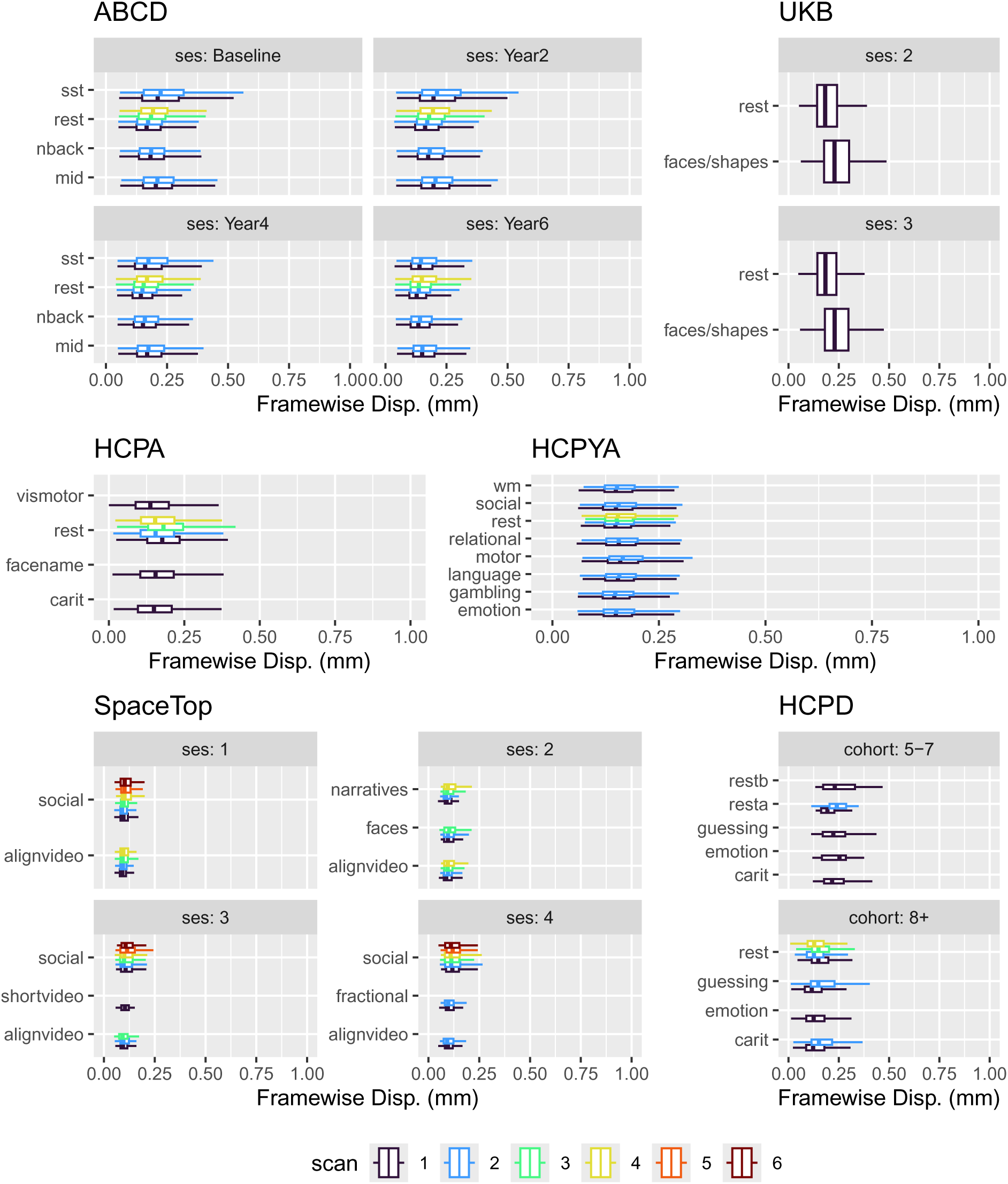
Average Motion Across all Conditions (Task and Rest). To facilitate comparisons across datasets, conditions are labeled according to their temporal order within a session (indicated by color). Panels correspond to sessions. Note that the three HCP studies are shown as having a single session, but conditions and scans were collected across days (e.g., in HCPYA, the first and second resting-state scans were collected on one day, and the third and fourth on another).

**Figure 2.**
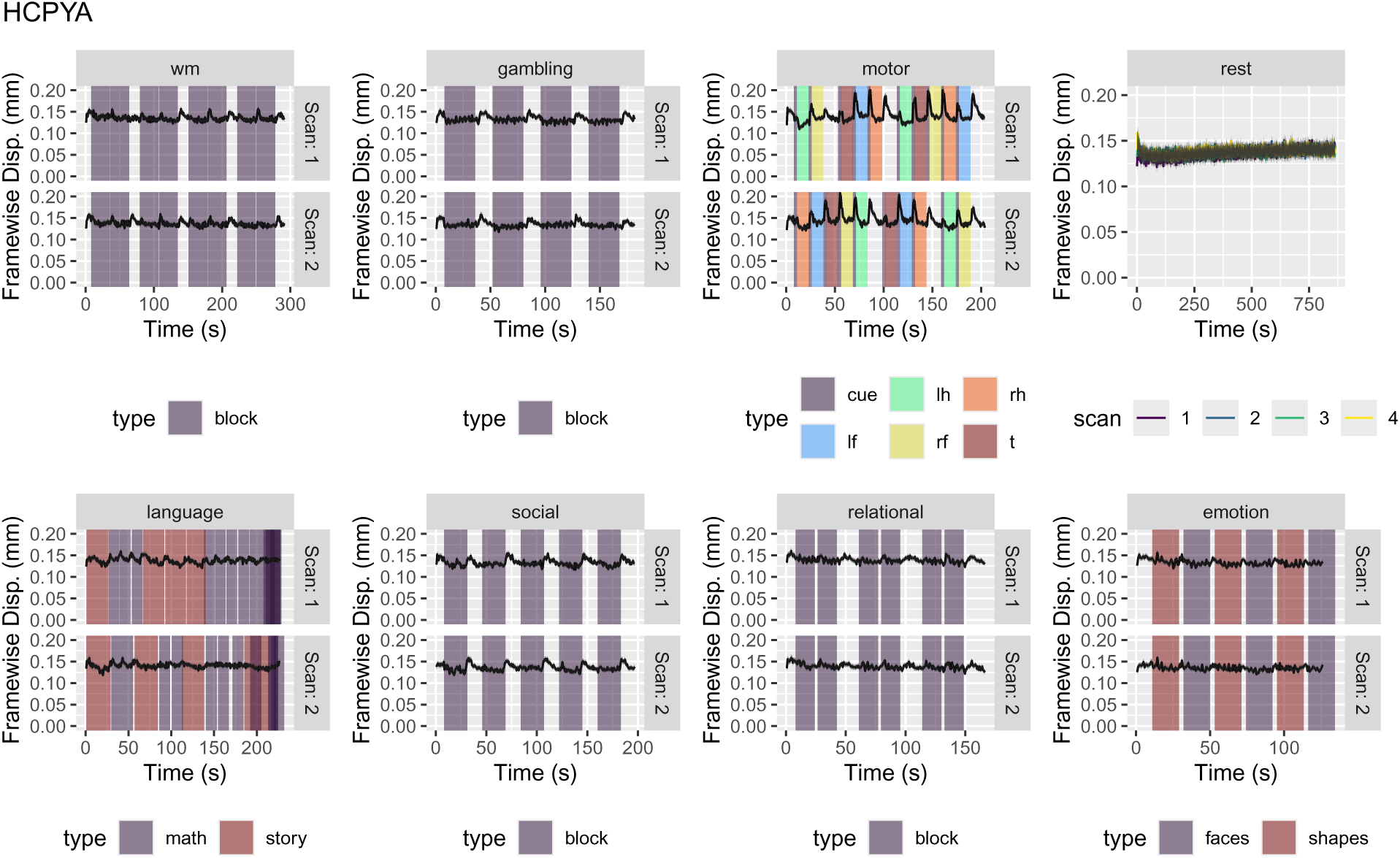
Average Motion by Time in the HCPYA Dataset. The lines trace the average framewise displacement (FD) across participants in the dataset, with ribbons marking 2x standard errors. Rectangular regions mark blocks of stimuli or responses (with timing averaged across participants).

Within many scans, motion exhibited three distinct temporal patterns (Figure 2 through Figure 7). **Settling-in effects**: In some datasets, motion in the first few frames was substantially lower than in subsequent frames (e.g., UKB, Figure 5), or conversely, higher (e.g., ABCD, Figure 6). While this could reflect different initial states of participant arousal or comfort, the differences may largely be driven by differences in reference volume for motion correction across datasets (e.g., the UKB uses a separate reference image, whereas the ABCD uses the temporal average volume). **Drift**: In most datasets, motion tended to increase throughout runs. Temporal changes in motion could be profound, with the average FD increasing by as much as 150 % (e.g., HCPA, CARIT task, Figure 3) from the start to the end of the scan. **Task-Correlated motion**: Motion was often highly correlated with the task structure. For example, the task scan in the UKB (Figure 5) used identical timing for all participants, producing visible blocks of elevated motion in the cross-participant average as well as sharp spikes at the moments of individual button presses. Similarly, the HCPYA Motor task (Figure 2) shows distinct motion profiles for different blocks (e.g., tongue movements vs. toe movements), illustrating how specific task demands physically perturb the head. These findings demonstrate that head motion is not a static property of a dataset but varies systematically across age, session position, and task structure. Such structured variation has important implications for how motion thresholds are applied and interpreted across large, heterogeneous cohorts.

**Figure 3.**
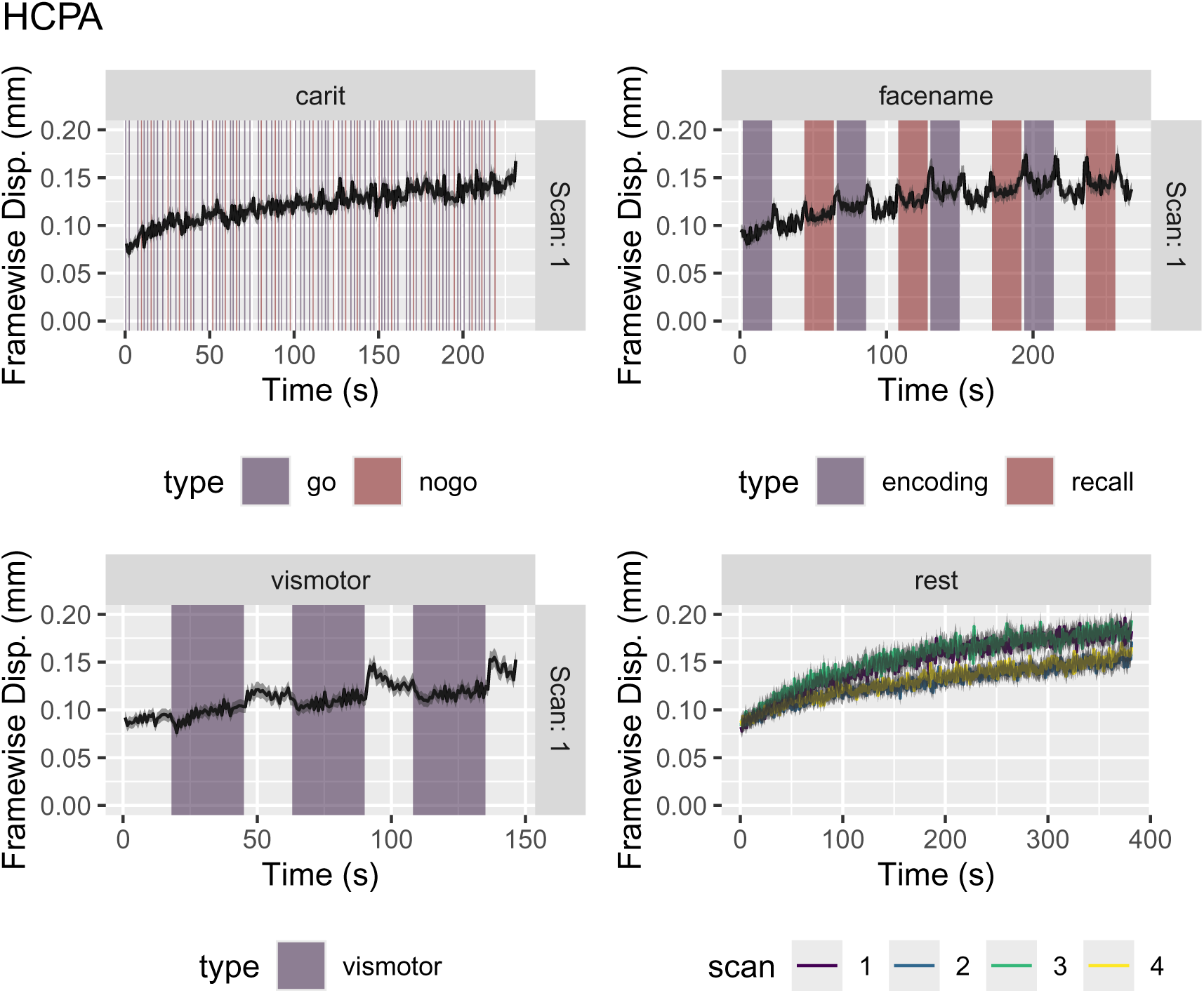
Average Motion by Time in the HCPA Dataset. Data plotted as in Figure 2.

**Figure 4.**
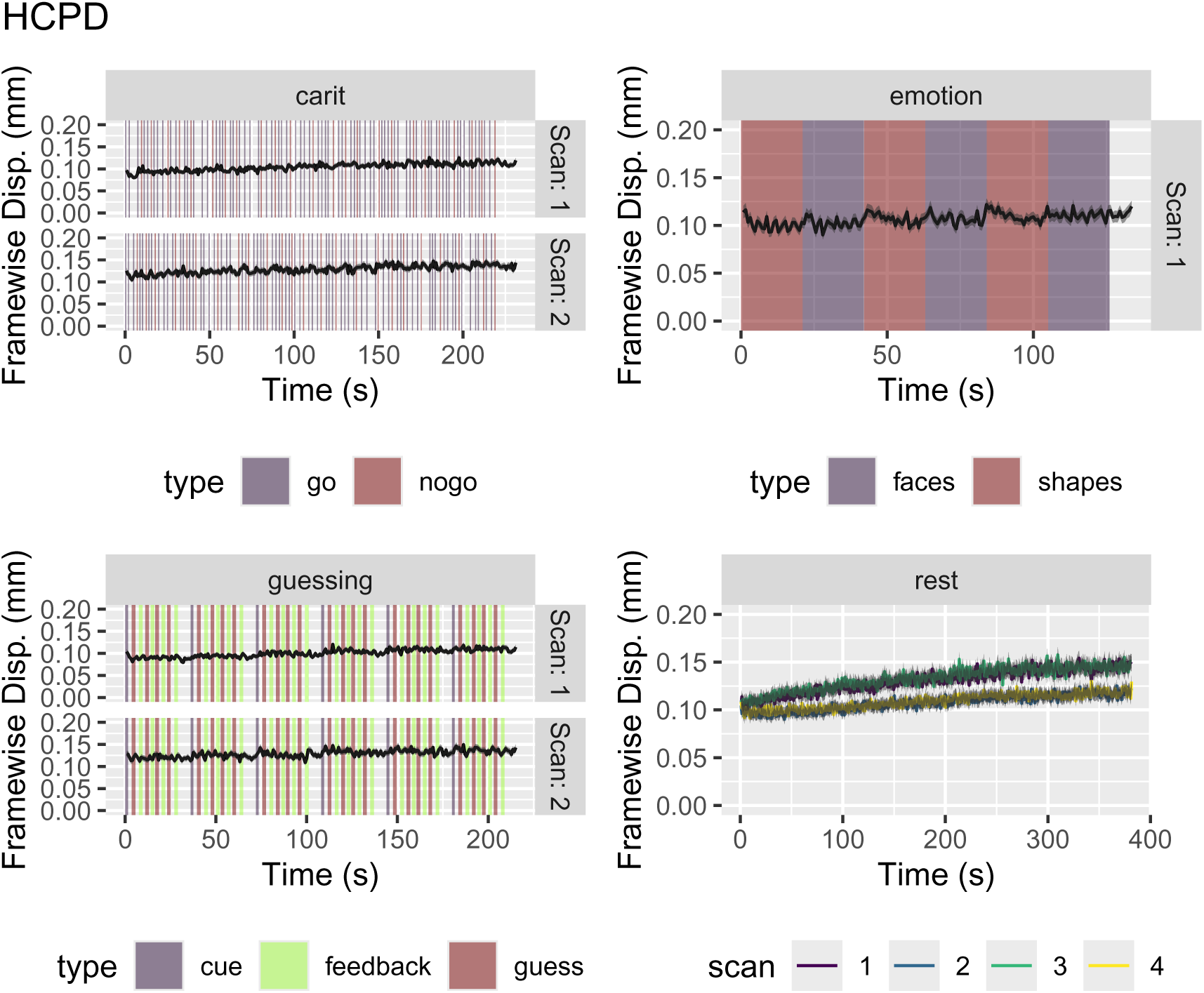
Average Motion by Time in the HCPD Dataset. Data plotted as in Figure 2.

**Figure 5.**
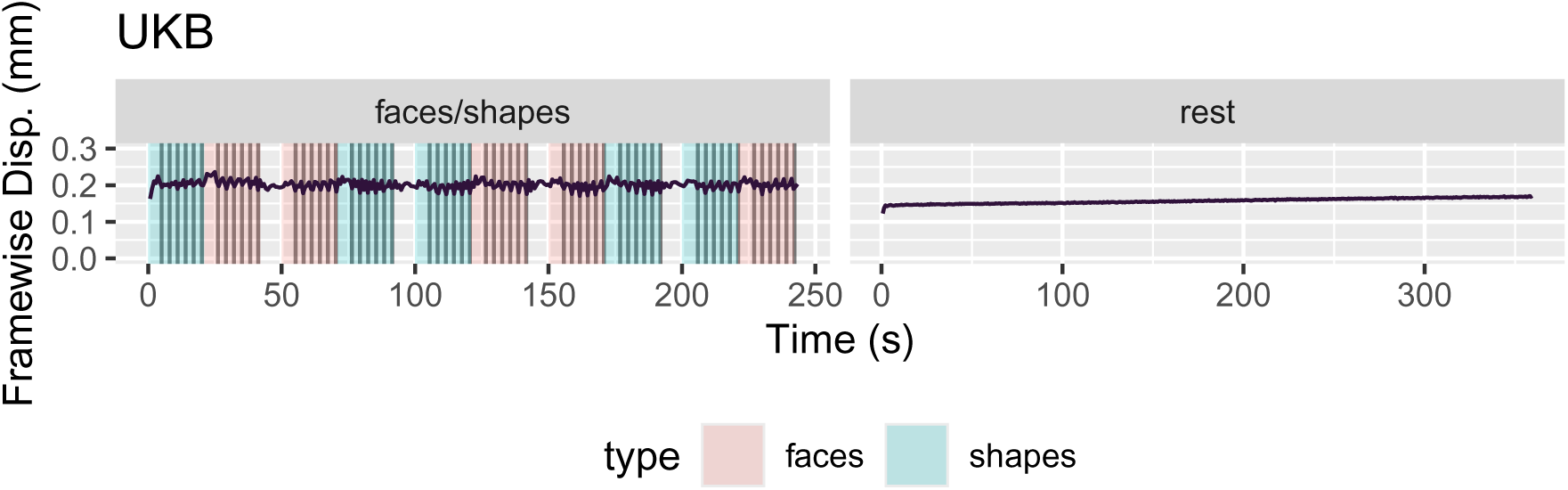
Average Motion by Time in the UKB Dataset. Data plotted as in Figure 2.

**Figure 6.**
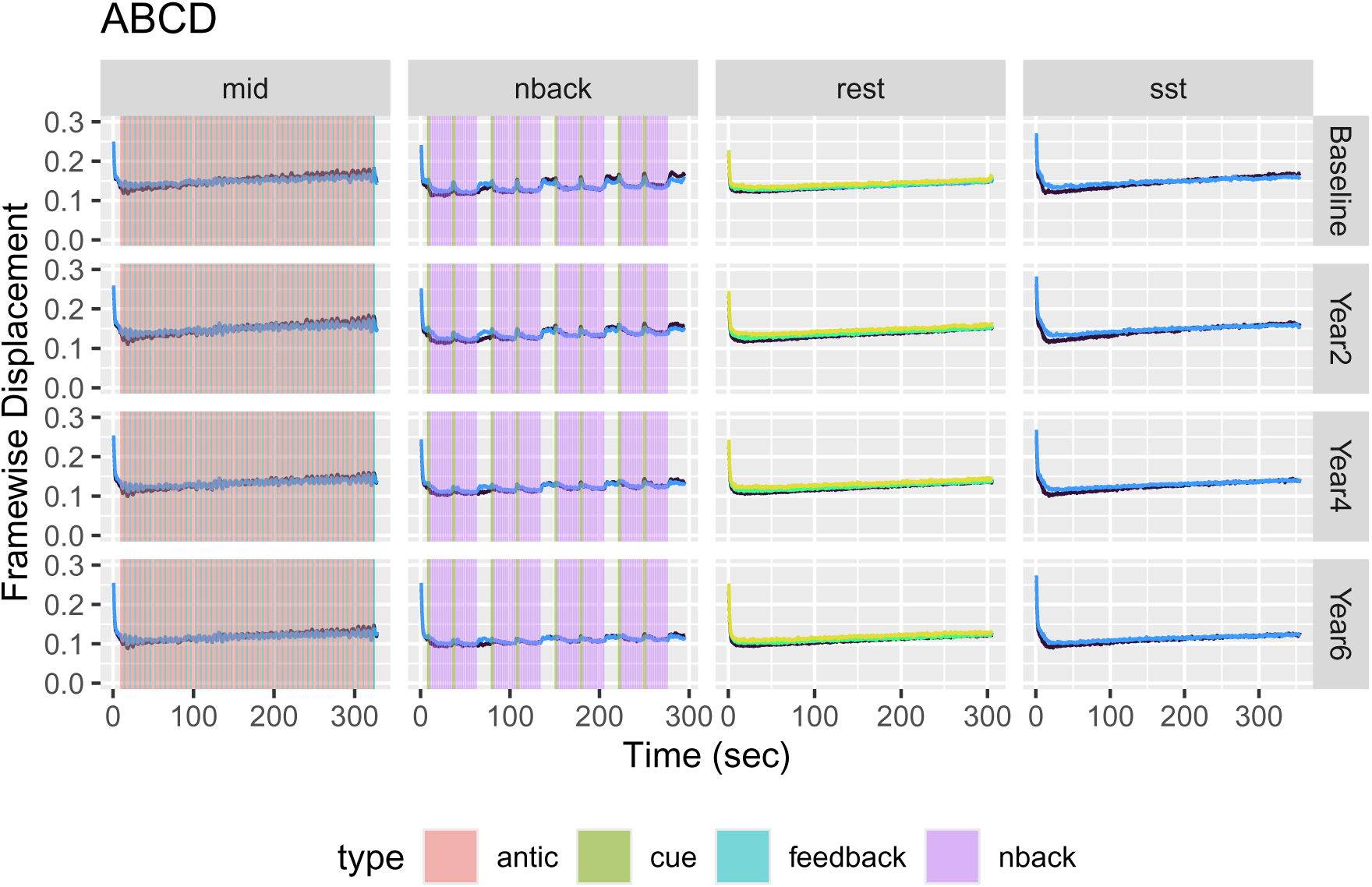
Average Motion by Time in the ABCD Dataset. Data plotted as in Figure 2.

**Figure 7.**
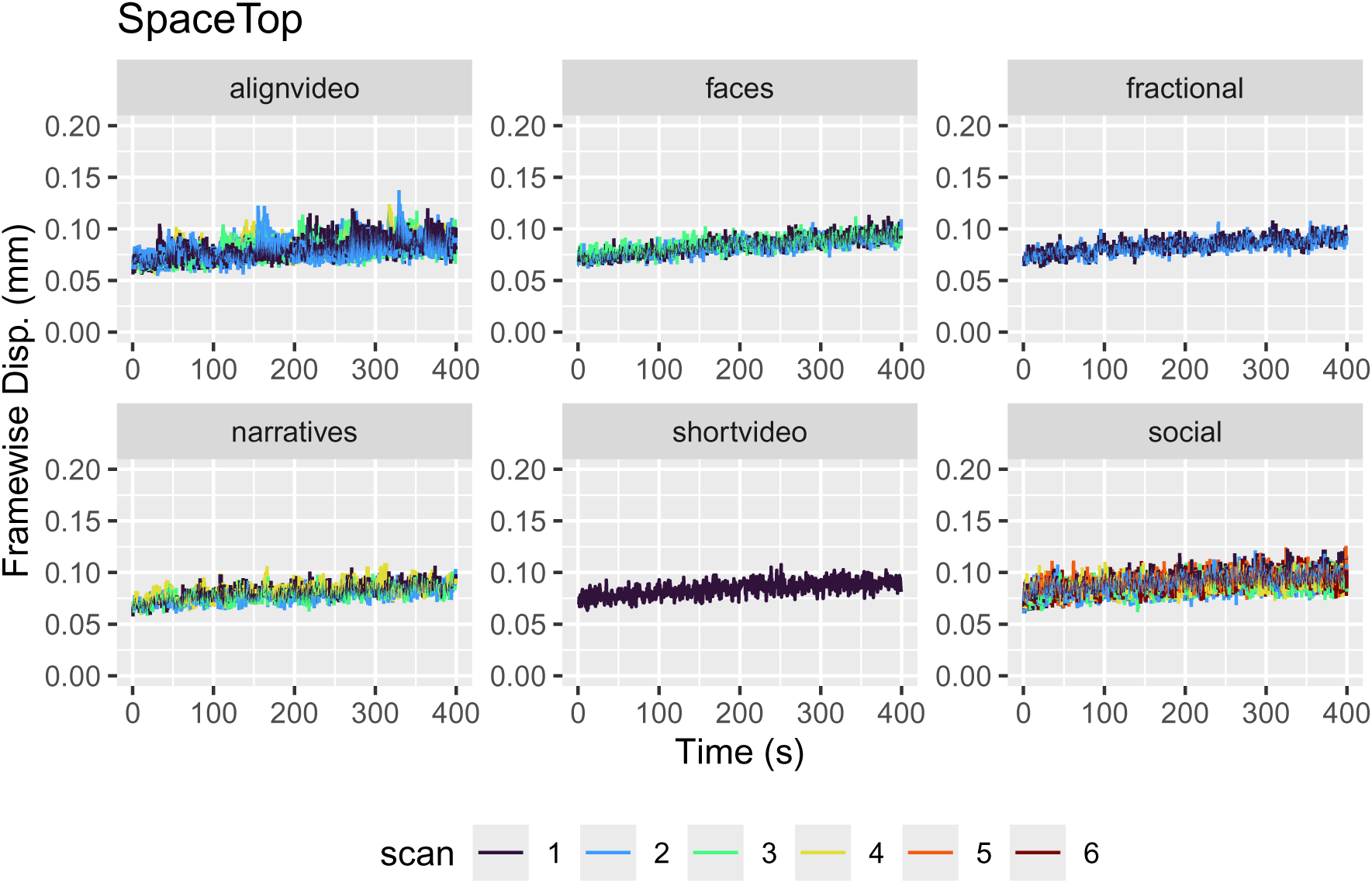
Average Motion by Time in the SpaceTop Dataset. Data plotted as in Figure 2.

### 3.2 Who Moves?

Across all datasets, there were 57084 unique individuals (76 % of whom were from UKB) and 320459 scans with motion data. The youngest participant was in the HCPD dataset (5.58 years old), and the oldest was in the HCPA dataset (over 85 years old). This broad age range enabled us to examine motion patterns across nearly the entire lifespan.

#### 3.2.1 Motion is Stable Across Scans and Sessions

To determine whether motion reflects transient noise or stable individual differences, we classified participants as “high-movers” or “low-movers” based on a single scan (Section 2.5) and compared motion between these groups across all remaining scans. Across datasets, individuals who exhibited high motion in one scan tended to exhibit high motion in all other scans. For example, participants in the HCPYA dataset who moved the most in the REST1 LR scan tended to move the most in all other scans (Figure 8 a). The separation between the high-mover (orange) and low-mover (blue) distributions was consistent across all seven tasks. Notably, within the ABCD dataset, groups defined based on motion in one scan from the baseline session had predictive power in follow-up sessions two, four, and six years later (Figure 8 b), even as head motion decreased at follow-up scans on average. In other words, ABCD participants who tended to move more at 9-10 years of age continued to move more throughout adolescence. This persistence across tasks and years indicates that head motion is not purely situational but reflects stable, trait-like characteristics of an individual, likely related to stable physiological or psychological factors (e.g., respiratory rate, motor control, anxiety).

**Figure 8.**
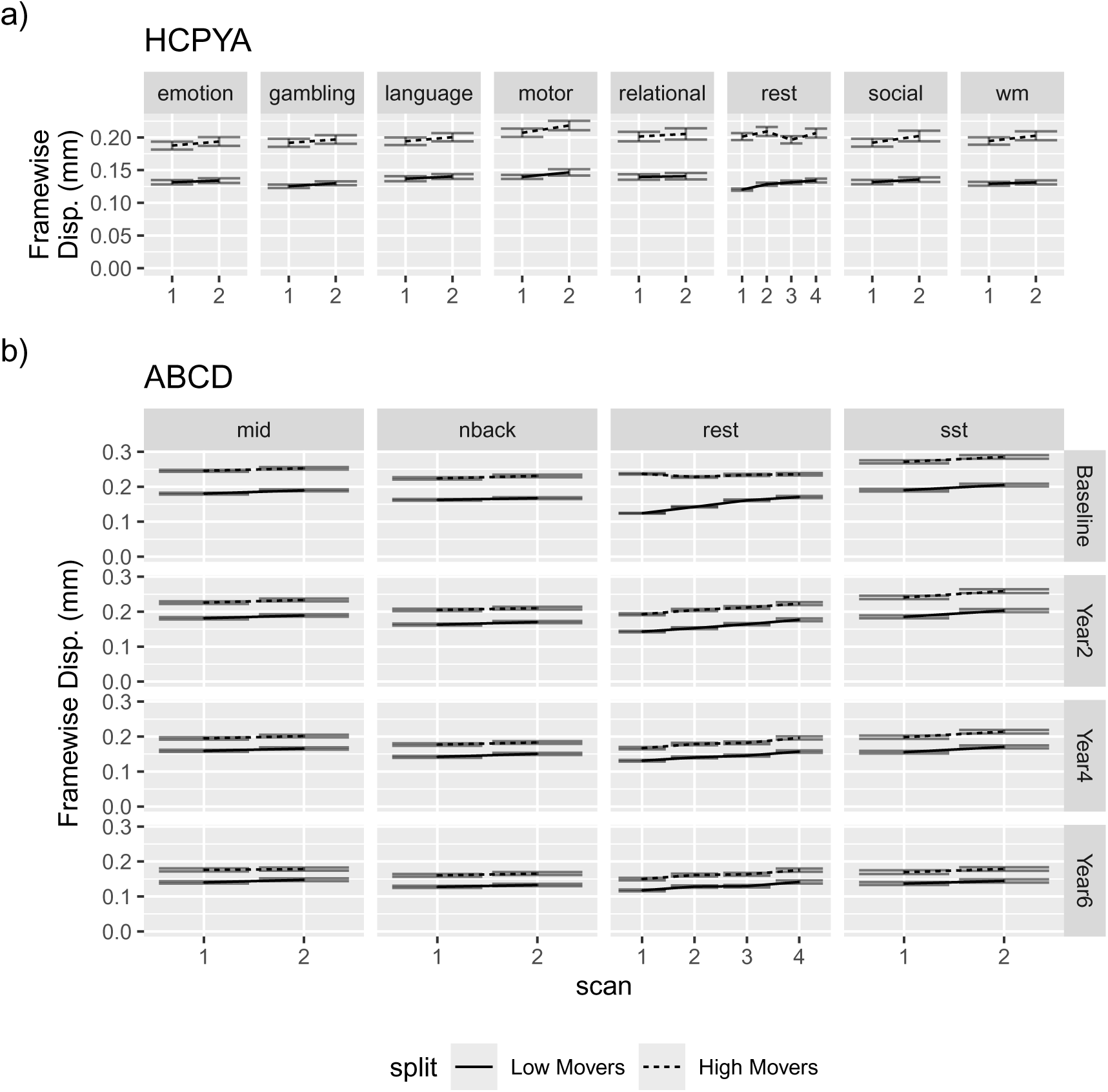
Motion is stable across scans. **(a)** High- and Low-Movers in the HCPYA dataset. Classification based on motion in REST1 generalizes to all other scans. **(b)** High- and Low-Movers in the ABCD dataset. Classification based on a single resting-state scan from the baseline session generalizes to sessions collected years later. MID: Monetary Incentive Delay; SST: Stop-Signal Task; NBack: Emotional N-back.

The HCP, UKB, and ABCD datasets include multiple resting-state scans, allowing us to measure consistency of mean motion within and across sessions. For context, we point to the categories provided by (Cicchetti & Sparrow, 1981): < 0.4: poor, [0.4, 0.6): fair, [0.6, 0.75): good, [0.75, 1): excellent.

We used the HCP and ABCD datasets to measure within-session consistency. In the HCP, consistency was _0.66_0.69_0.72_ (*F*(693, 2082) = 10.02, *p* < 0.001), _0.56_0.6_0.63_ (*F*(608, 1827) = 6.88, *p* < 0.001), and _0.77_0.79_0.81_ (*F*(875, 2628) = 15.79, *p* < 0.001), in the HCPA, HCPD, and HCPYA, respectively. Likewise, we calculated the consistency of motion in the resting-state scans within each session of ABCD, finding values of _0.55_0.57_0.58_ (*F*(3338, 10017) = 6.22, *p* < 0.001) at baseline, _0.57_0.58_0.6_ (*F*(3220, 9663) = 6.58, *p* < 0.001) in Year 2, _0.61_0.62_0.64_ (*F*(3148, 9447) = 7.62, *p* < 0.001) in Year 4, and _0.64_0.66_0.67_ (*F*(2424, 7275) = 8.71, *p* < 0.001) in Year 6.

With the UKB and ABCD, we measured reliability across sessions. In the UKB, the consistency for average FD in the resting-state scan is _0.73_0.73_0.74_ (*F*(43332, 43333) = 6.44, *p* < 0.001), and in the ABCD it was _0.4_0.41_0.42_ (*F*(10742, 32229) = 3.76, *p* < 0.001).

Overall, the amount of motion in these datasets was comparable to that observed in the MRIQC aggregation of over 1.5 million scans (compare Figure 9 d to Figure 9 a-c). Crucially, because motion is both trait-like and demographically structured, exclusion based on motion is unlikely to operate randomly across participants. Instead, it preferentially removes individuals drawn from specific age groups, body compositions, and, to a lesser extent, sex categories.

**Figure 9.**
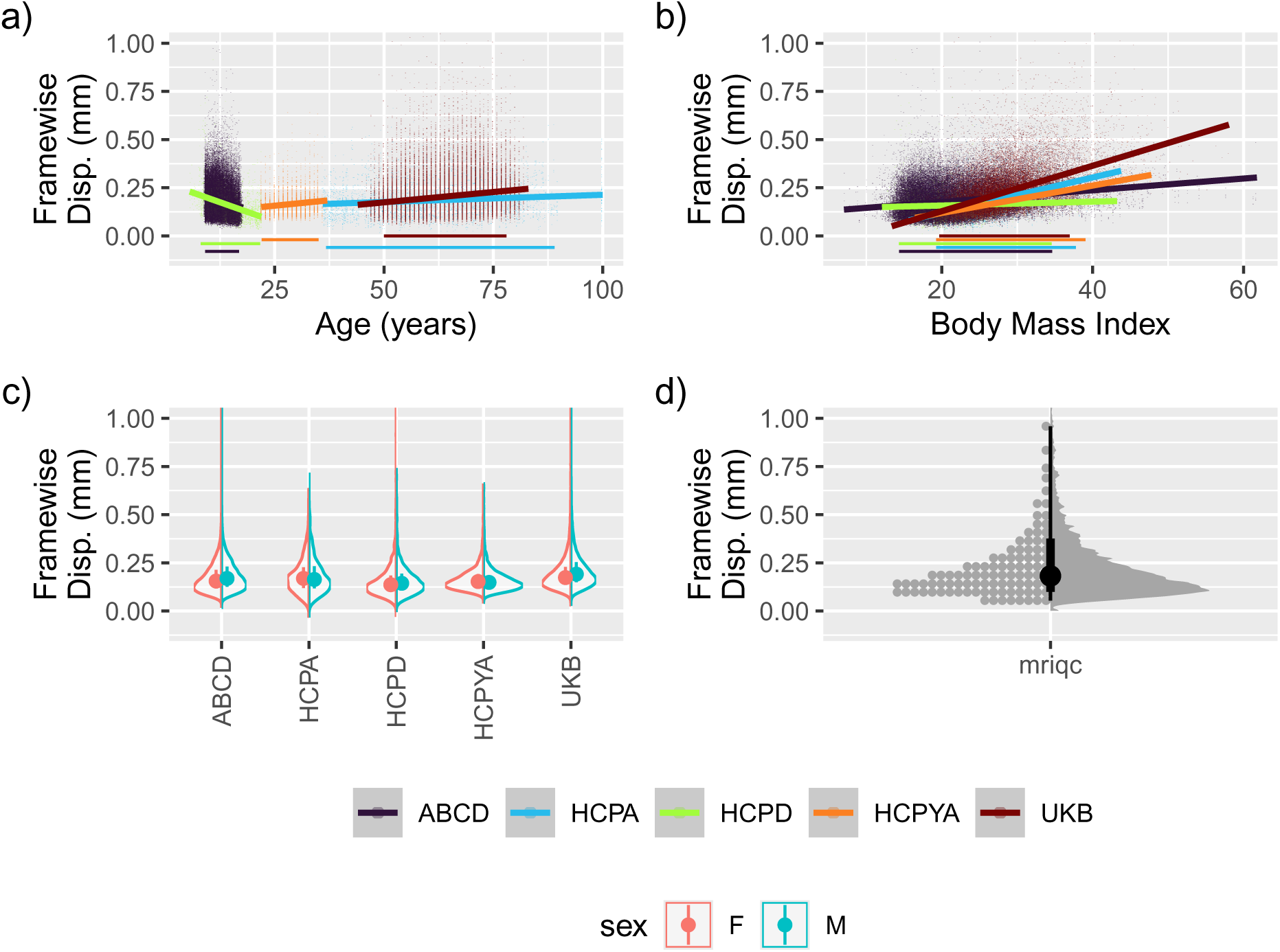
Average Motion in Resting-State Scans by **(a)** age, **(b)** body-mass index (BMI), and **(c)** Sex. For reference, the average framewise displacement of the MRIQC dataset is shown in **(d)**. For the MRIQC dataset, each point corresponds to a percentile (the highest percentile is not shown), and the slabs correspond to the 0.66- and 0.95-quantile intervals.

#### 3.2.2 Demographic Patterns of Motion

Having established that motion is stable within individuals, we next examined how motion varies across participant characteristics known to be associated with movement: age, body mass index (BMI), and sex (Figure 9). The variables are summarized for the high- and low-motion subgroups in Table 3.

**Table 2.**
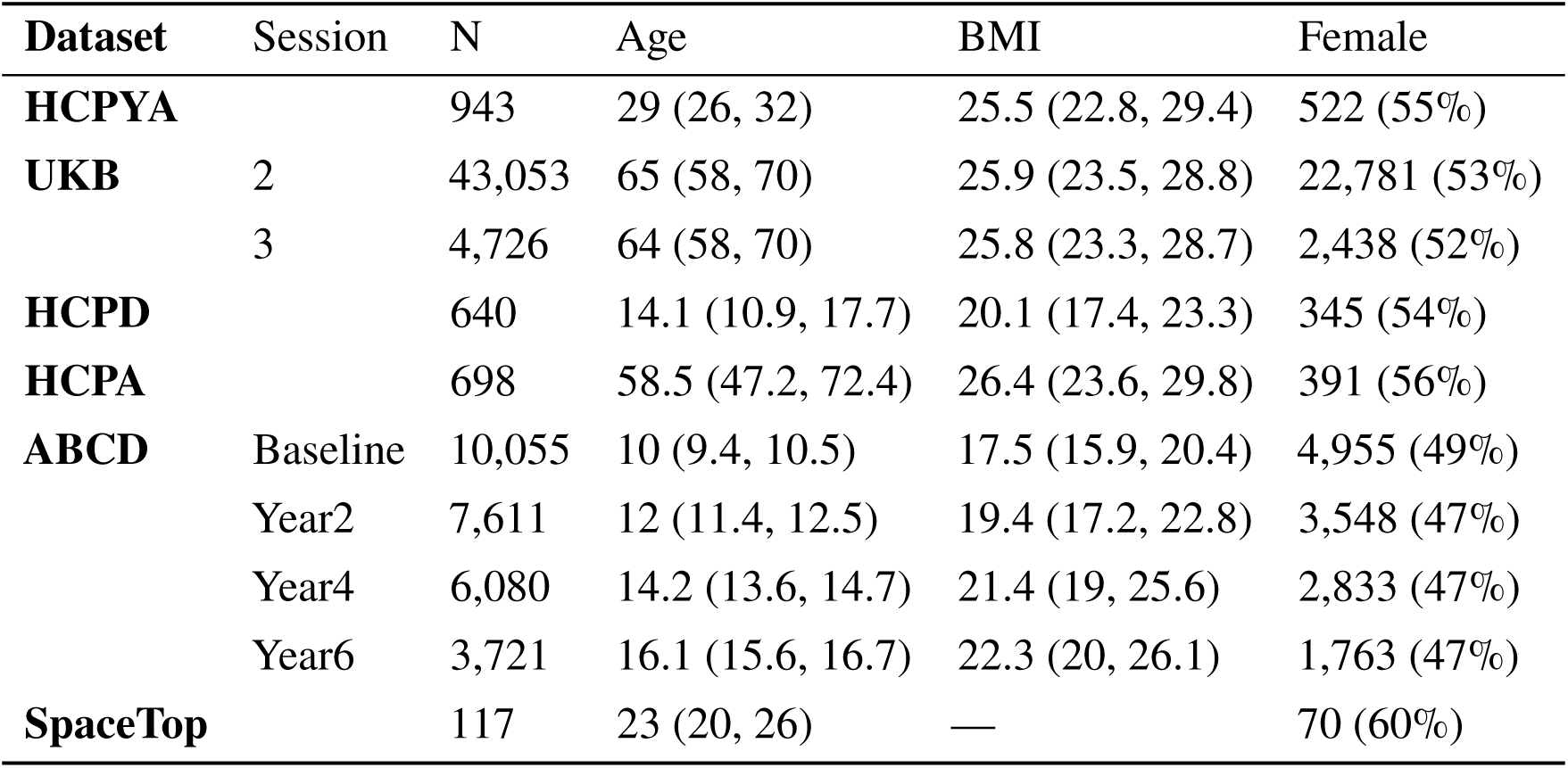
Participant Demographics. Age and BMI are shown as median (Q1, Q3); Female as n (%). BMI was not recorded for SpaceTop. Sessions are listed only for datasets with repeated measures.

**Table 3.** Demographic Variables in High and Low Motion Subgroups (groups defined as in Figure 8). Age and BMI are shown as median (Q1, Q3); sex as n (%). In the ABCD columns, demographics refer to the baseline session.

| Characteristic | ABCD (Baseline) |  | HCPYA |  |
| --- | --- | --- | --- | --- |
|  | Low | High | Low | High |
| N | 4,103 | 4,102 | 471 | 470 |
| Female | 2,158 (53%) | 1,991 (49%) | 248 (53%) | 273 (58%) |
| Male | 1,945 (47%) | 2,111 (51%) | 223 (47%) | 197 (42%) |
| Age | 10.1 (9.5, 10.6) | 9.9 (9.4, 10.5) | 29 (26, 31) | 29 (26, 32) |
| BMI | 16.9 (15.6, 18.8) | 18.3 (16.2, 21.5) | 23.6 (21.6, 26) | 28.4 (25.2, 32.4) |

##### Age

Average framewise displacement exhibited a U-shaped relationship with age (Figure 9 a), with the youngest (HCPD, ABCD) and oldest (HCPA, UKB) participants moving more than young and middle-aged adults. This pattern is consistent with developmental differences in motor control and age-related changes in physiology or comfort during scanning. However, substantial differences were also observed across datasets with overlapping age ranges. For example, adolescents in ABCD exhibited higher motion than similarly aged participants in HCPD. To test this quantitatively, we fitted a linear mixed model to predict average motion and contrasted the two datasets (Section A.3). According to that analysis, a 9-year-old participating in the ABCD or HCPD would display a statistically significant difference in motion (_−0.021_−0.014_−0.007_, *p* < 0.001), indicating that motion is lower by around 0.01 mm in the HCPD dataset.

##### BMI

Replicating previous findings (Bolton et al., 2020; Teo et al., 2024; Ward et al., 2024), we observed that motion increased with BMI across datasets (Figure 9 b). In resting-state scans, the effect of BMI was significant (_0.002_0.003_0.003_, *p* < 0.001). Despite its small effect size at the individual level, this association has population-level implications in large cohorts where BMI is unevenly distributed.

##### Sex

There was a statistically significant difference for sex, with females moving slightly less than males (estimated marginal mean contrast at average age 44.9 and average BMI 24.45: _−0.008_−0.007_−0.006_, *p* < 0.001). Importantly, sex differences were more pronounced in younger cohorts (Figure 9 c), suggesting developmental interactions. Indeed, at the BMI that is equal to the average for ABCD (20.95), there was a significant interaction between the effect of sex at age 9 as compared to the effect of sex at age 15 (_−0.004_−0.004_−0.003_, *p* < 0.001), indicating that sex differences diminish with increasing age during development.

#### 3.2.3 Motion as a Function of Psychological Characteristics

Next, we selected one dataset with a rich collection of curated psychological variables, the HCPYA, and assessed how each relates to motion separately (Figure 10). We fit an ordinary least-squares linear model predicting the psychological variable from motion. Several of the assessed variables were positively associated with motion, with higher values indicating greater motion. One variable, avoidance, was negatively associated with motion: more avoidant individuals were less likely to move.

**Figure 10.**
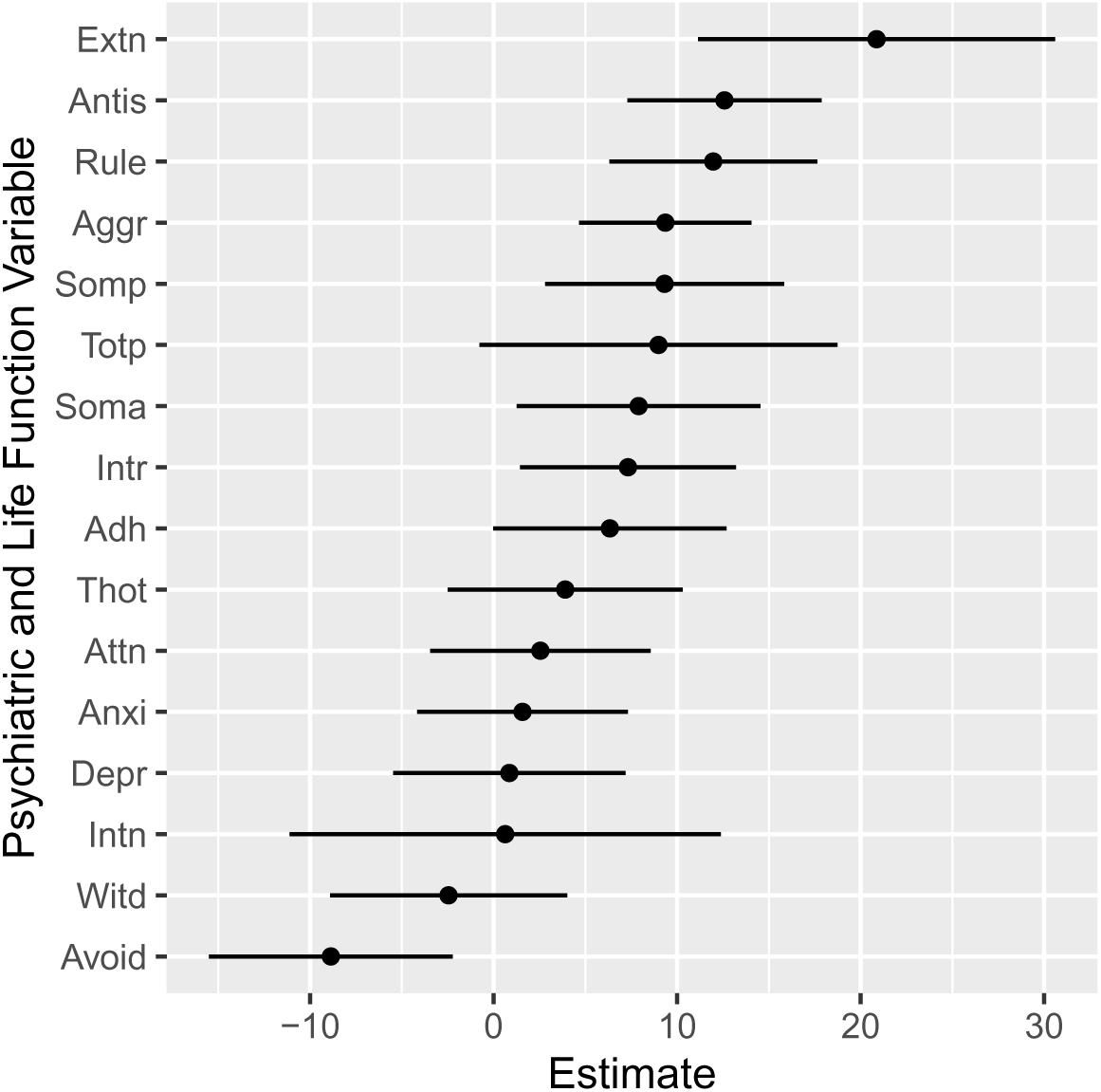
Relationships Between Average Motion and Psychological and Life Function Variables within HCPYA. All measures are standardized for age and sex, and the reported estimates are from univariate analyses. Lines show 95 % confidence intervals. Extn: Externalizing; Antis: Antisocial Personality Problems; Rule: Rule Breaking Behavior; Soma: Somatic Problems; Aggr: Aggressive Behavior; Totp: Achenbach Adult Self-Report Total; Intr: Intrusive Thoughts; Soma: Somatic Complaints; Adh: AD/H Problems; Thot: Thought Problems; Attn: Attention Problems; Anxi: Anxiety Problems; Depr: Depressive Problems; Intn: Internalizing; Witd: Withdrawn; Avoid: Avoidant Personality Problems.

### 3.3 Who is Excluded?

To quantify the practical consequences of motion-based quality control, we simulated participant exclusion under two commonly used regimes: a “lenient” and a “strict” threshold (Section 2.3). Across datasets, motion-related exclusion rates were substantial and highly variable, indicating that identical thresholds can produce dramatically different levels of data loss depending on cohort characteristics (Figure 11).

**Figure 11.**
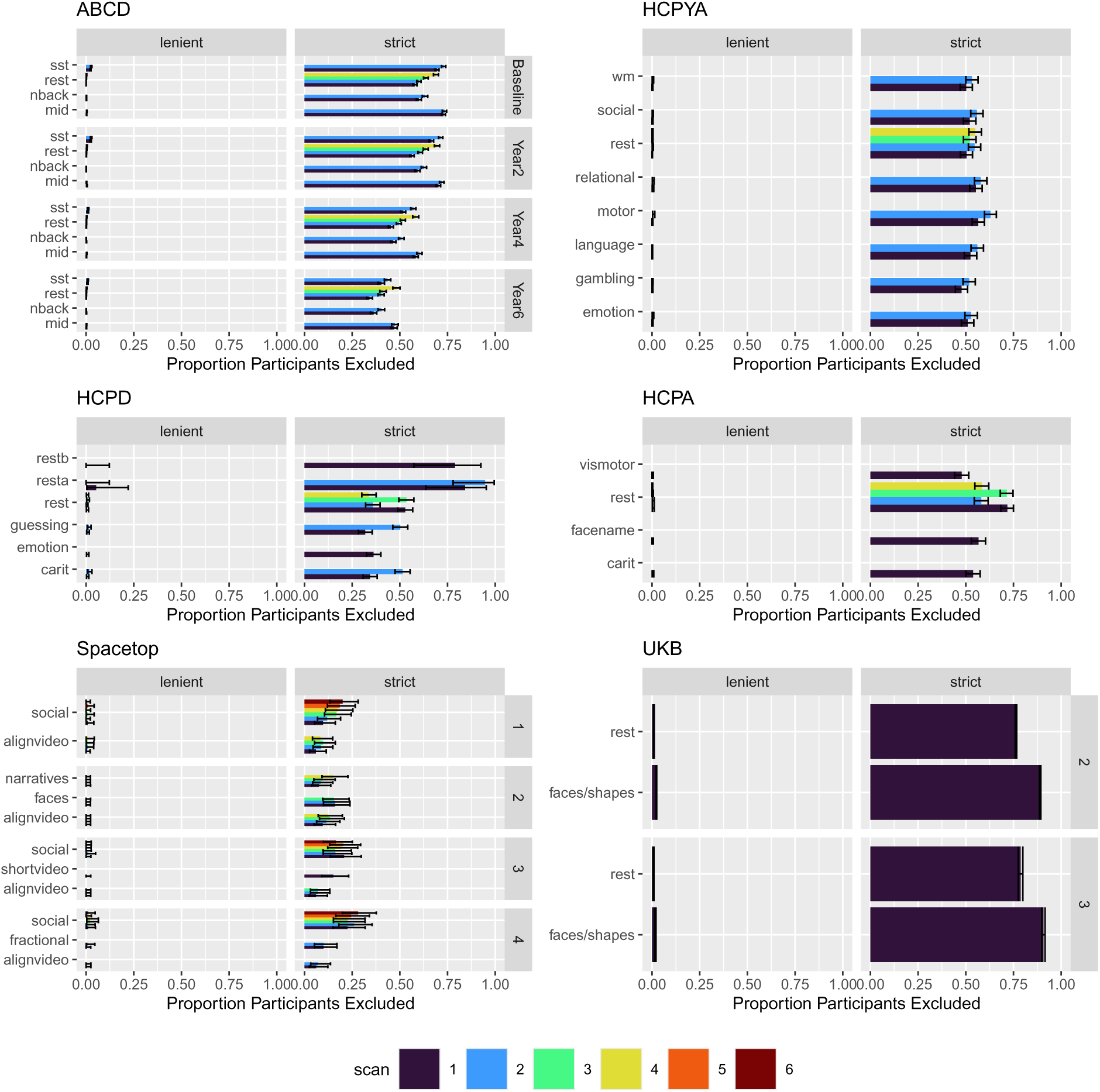
Rates of Participant Exclusion by Framewise Displacement. Proportions were calculated within each scan type (e.g., each combination of session, run, and task). Lenient and Strict refer to exclusion regimes (see Methods). The variable “scan” (color) refers to the order in which scans were collected (including scans collected on separate days). See also Figure 12.

Under the strict regime, more than half of the participants would be excluded from several datasets. In the UKB task data, strict thresholds would exclude over 80 % of participants, substantially reducing the usable sample size. In contrast, fewer than 10 % of participants would be excluded in the SpaceTop dataset, highlighting the exceptionally high data quality of this smaller, intensively sampled cohort. The ABCD dataset exhibited particularly high baseline exclusion rates, consistent with the high motion observed in younger participants.

These results demonstrate that motion thresholds do not operate uniformly across studies. Instead, they interact with the demographic and behavioral composition of each cohort to determine the effective analytic sample.

#### 3.3.1 Impact of Respiratory Filtering

Because respiratory-related “pseudo-motion” may inflate framewise displacement in adults, we repeated all exclusion analyses using notch-filtered motion parameters (Section 2.4). Removing frequencies associated with respiration reduced exclusion rates across datasets (compare Figure 11 to Figure 12), particularly in adult cohorts. This suggests that a portion of exclusion in adult populations may be driven by physiological artifacts rather than mechanical head movement, and that preprocessing decisions can meaningfully alter participant retention.

**Figure 12.**
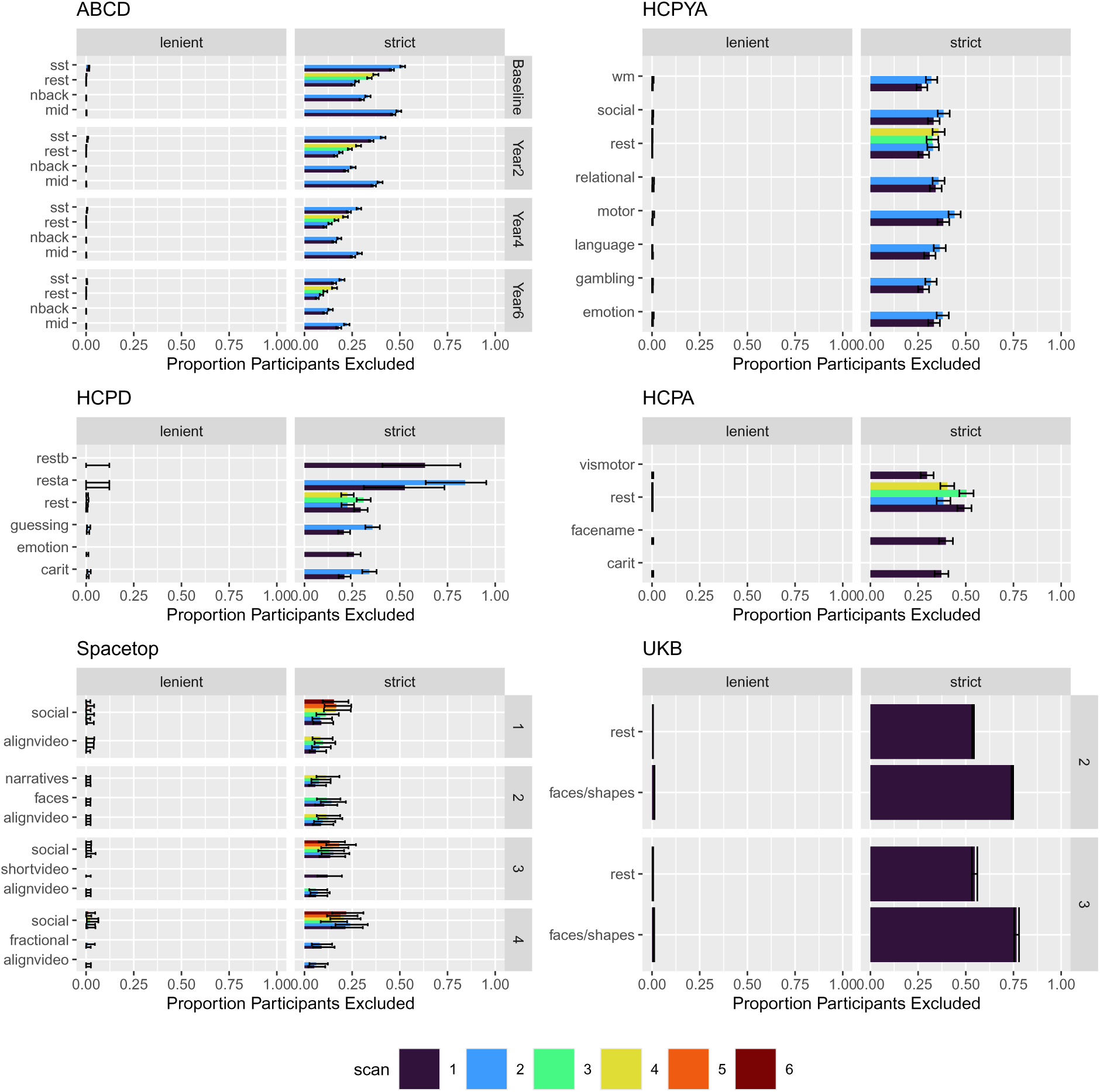
Rates of Participant Exclusion by Filtered Framewise Displacement. Data are plotted as in Figure 11.

#### 3.3.2 Demographic Patterns of Exclusion

We next examined how exclusion rates varied across demographic categories. Figure 13 displays the proportion of participants lost to thresholding as a function of Age, BMI, and Sex.

**Figure 13.**
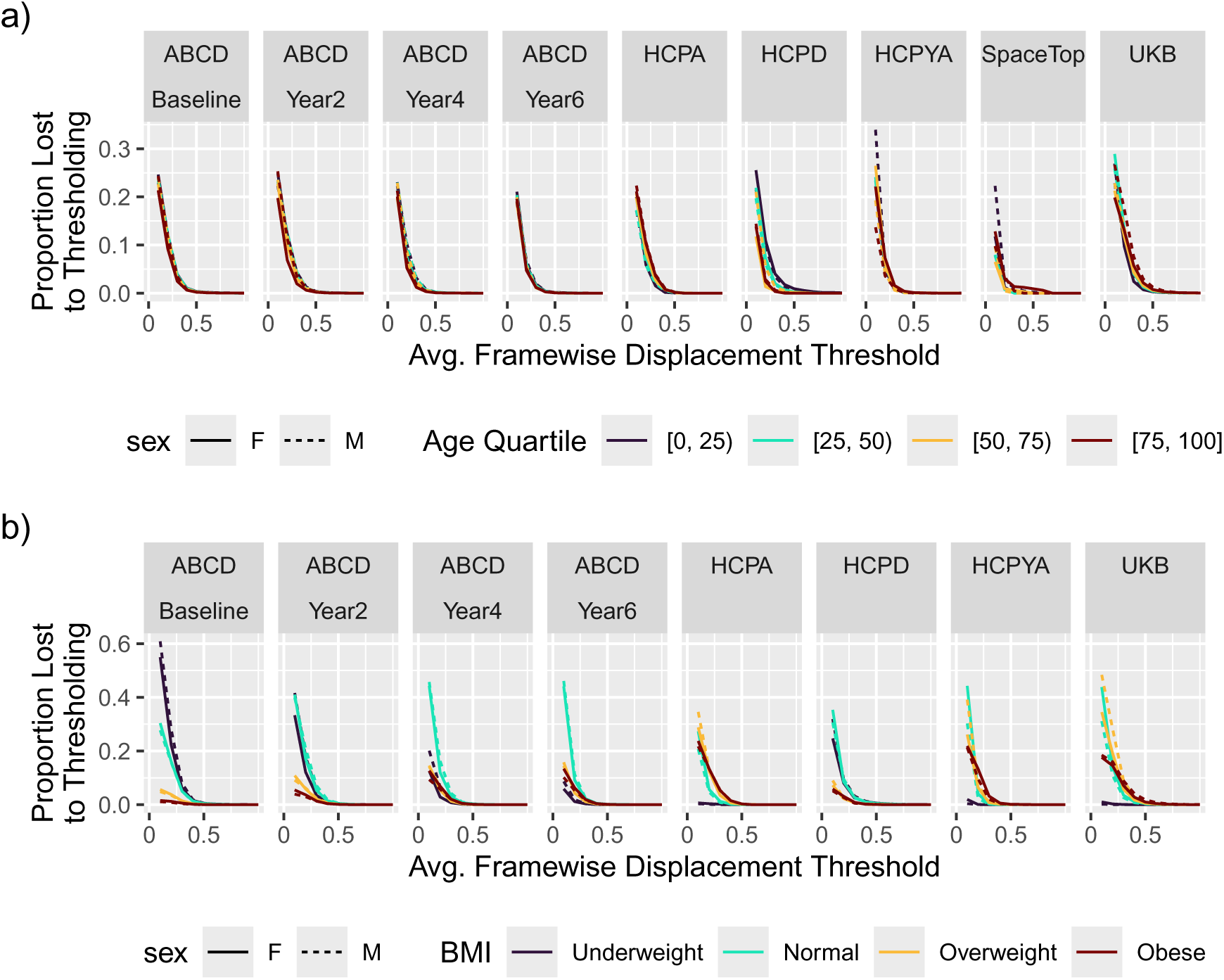
Rates of exclusion by **(a)** Age, **(b)** BMI, and **(c)** Sex.

##### Age

Strict exclusion criteria disproportionately excluded both the youngest and oldest participants. In ABCD and HCPD, children were excluded at markedly higher rates than young adults. In UKB and HCPA, the “oldest-old” exhibited higher exclusion probabilities than middle-aged adults. Thus, motion-based exclusion amplifies the U-shaped age distribution observed in raw motion data, selectively reducing representation at both ends of the lifespan.

##### BMI

Exclusion rates increased monotonically with BMI. In the UK Biobank, participants in the “Obese” category were excluded at nearly twice the rate of those in the “Normal” BMI category under strict thresholds. Because BMI is associated with a wide range of health outcomes, this disproportionate exclusion introduces the potential for systematic distortion (collider bias) in studies examining metabolic, inflammatory, or cardiovascular correlates of brain function.

##### Sex

Exclusion rates were relatively similar between sexes, though males in the ABCD cohort were excluded at slightly higher rates than females, consistent with the observed sex difference in raw motion. Although sex-based exclusion differences were modest relative to age and BMI, they further illustrate that motion thresholds do not operate independently of participant characteristics.

### 3.4 What are the Impacts of Exclusions?

Finally, we turn to the impact of motion on downstream analyses, focusing on the biases introduced by either retaining motion-corrupted volumes or excluding high-motion participants. First, we asked whether motion in these datasets influences downstream functional-connectivity estimates even after modern preprocessing (e.g., Power et al., 2012). Modern datasets often include sophisticated data cleaning, in part to account for the high degree of noise expected in community samples. One possibility is that these procedures mitigate motion-related bias sufficiently to reduce the need for aggressive participant exclusion.

To address this, we selected a subset of 426 participants from the UKB (average resting-state motion in the first session: 0.2 mm, standard deviation: 0.09) and calculated the mean absolute change (MAC; Section 2.7.1). We note, however, that MAC provides only an indirect assessment of preprocessing sufficiency, in that a higher MAC may reflect reduced bias, increased estimate variability due to frame removal, or both. Nevertheless, several patterns are informative. The overall higher MAC for BP+smooth data indicates a larger effect of censoring in the uncleaned data (Figure 14), consistent with preprocessing mitigating some noise. For both cleaning strategies considered, censoring by DPD has a larger effect than censoring by framewise displacement at more stringent thresholds (compare red vs. blue lines in Figure 14). Assuming that DVARS is a more accurate measure of data quality than framewise displacement (even when the level of quality is driven by motion, e.g., Phạm et al. (2023)), then the higher MAC observed after DPD-based censoring suggests that residual bias remains in the functional connectivity estimates after both preprocessing strategies. Put differently, functional connectivity estimated after these preprocessing strategies may still be adversely affected by motion-corrupted volumes. Having shown results consistent with a lingering impact of motion on preprocessed data, we next examined the effects of excluding participants from the studies. We first examined how observed exclusion rates affect the statistical power to detect a brain-wide association in datasets with a resting-state scan (e.g., a correlation between regional volume and a characteristic of interest, such as depression status). For reference, note that a common correlation for these associations is 0.1 (Marek et al., 2022). As seen in Figure 16, across all datasets, the lenient motion threshold produced little reduction in power. The strict threshold had a larger impact, although the impact was only substantial in the smallest datasets (*n* < 1000). That is, with a sufficiently large dataset (e.g., UKB, ABCD), excluding many people still results in high power. In HCPA, assuming an effect size of 0.1, power dropped from 0.754 to 0.288 when excluding participants based on raw motion traces, and to 0.469 when using filtered traces. HCPYA straddled the conventional target of 0.8 power, dropping from 0.867 in the full dataset to 0.581 or 0.742, depending on the exclusion rule (strict not filtered, strict filtered).

**Figure 14.**
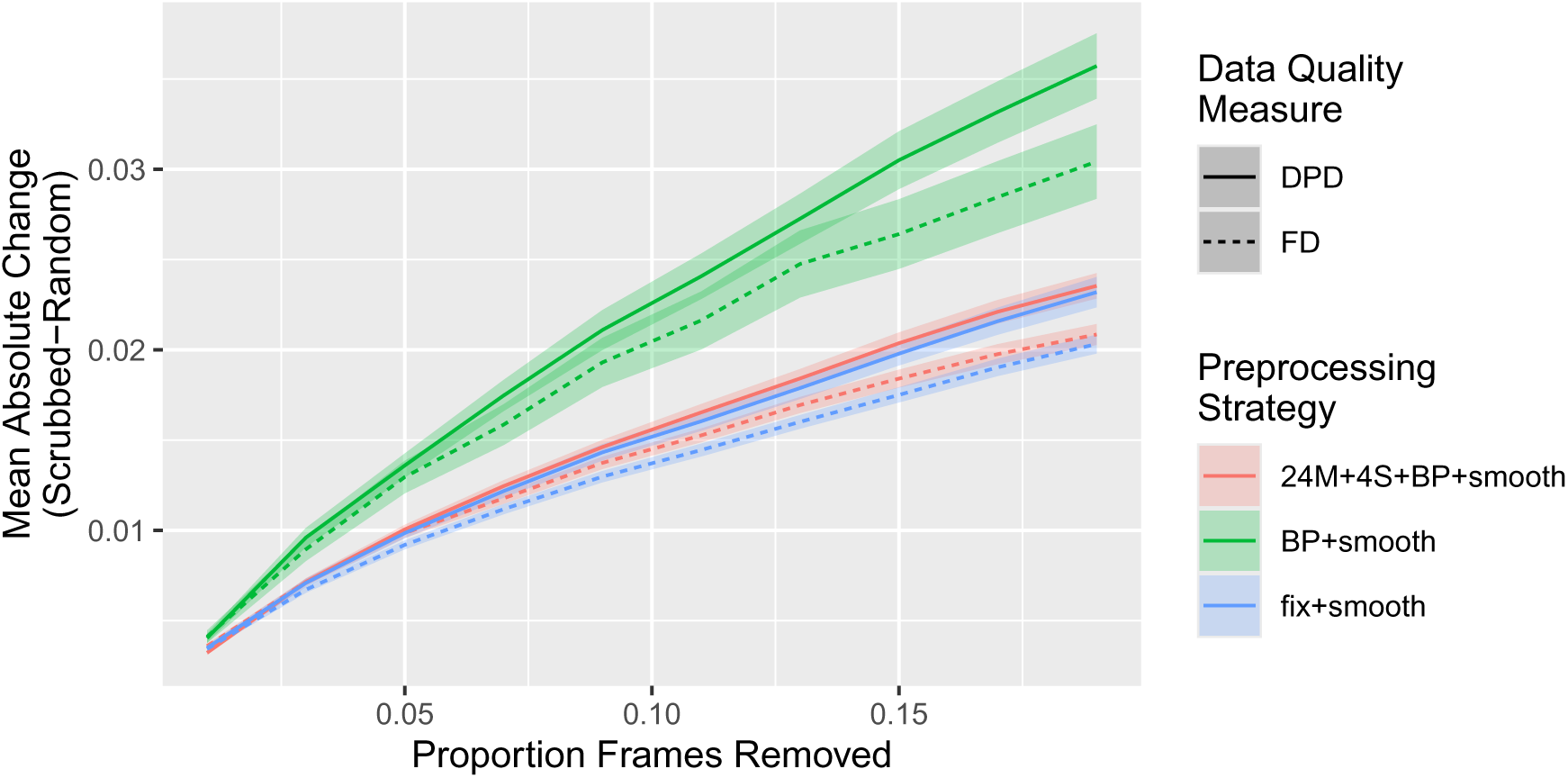
Mean Absolute Change in Functional Connectivity. Lines trace the average (across participants), and the ribbons span two standard errors. Higher values indicate a larger (beneficial) effect of censoring, but comparisons are only meaningful for a given censoring proportion. Colors indicate preprocessing strategy. 24M: 24 motion regressors (3 translation, 3 rotation, squares, first-order differences, and squares of first-order differences); 4S: 4 spatial parameters (CSF, WM, first-order differences); BP: bandpass filter; smooth: Gaussian Smoothing; fix: standard UKB independent-component analysis-based cleaning; DPD: change in percent DVARS within the uncleaned data; FD: framewise displacement.

**Figure 15.**
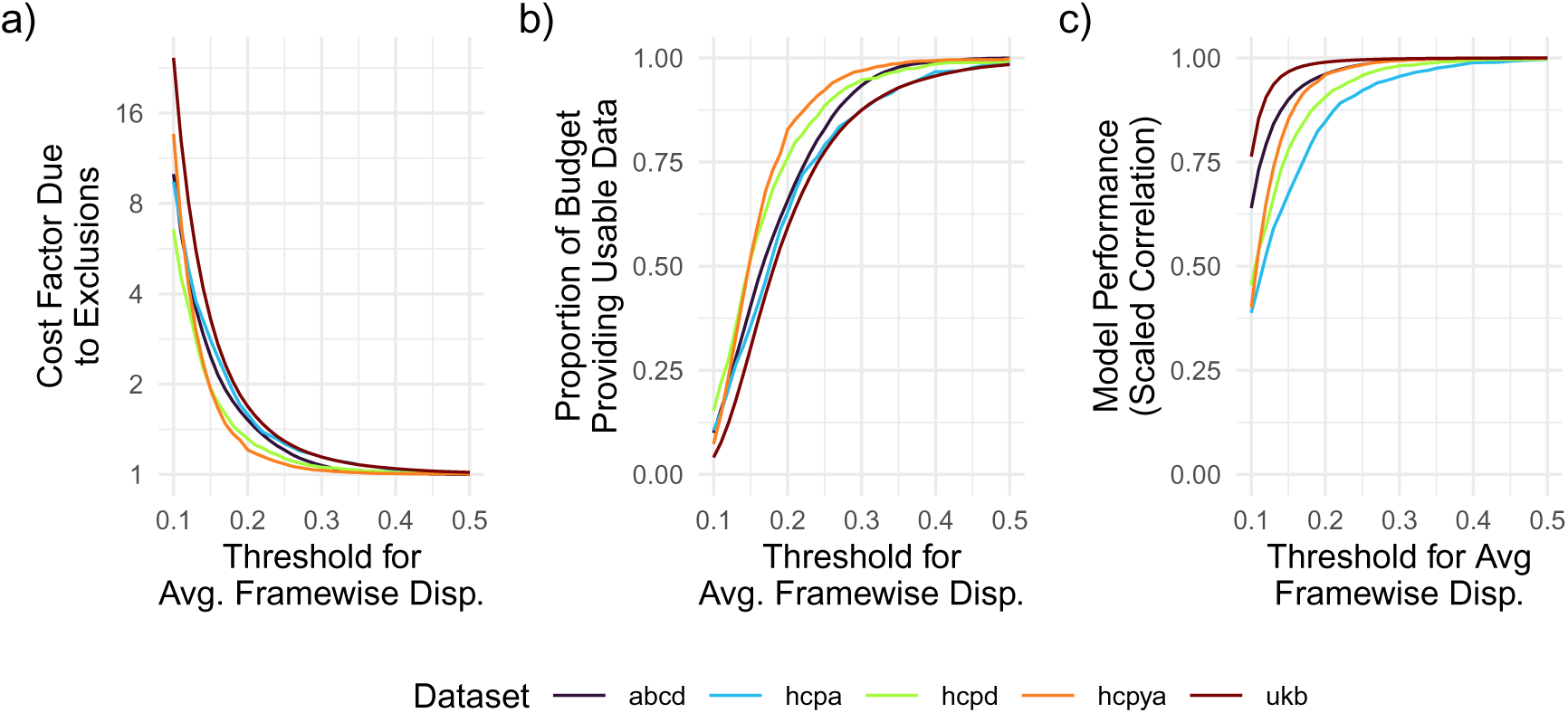
Consequences of Motion for Data Usability and Model Performance. (**(a)**) Estimated cost inflation when accounting for motion-related exclusions, assuming sampling continues until the target size is reached. For each dataset, the target size equals the full sample size. (**(b)**) Fraction of the total budget resulting in usable data under fixed budgets. (**(c)**) Reduction in predictive performance under fixed budgets when participants are excluded due to motion. Maximum performance is based on using the complete dataset. In (**(a)**)-(**(c)**), exclusion rates are based on the proportion of participants that would be excluded given the observed motion levels.

**Figure 16.**
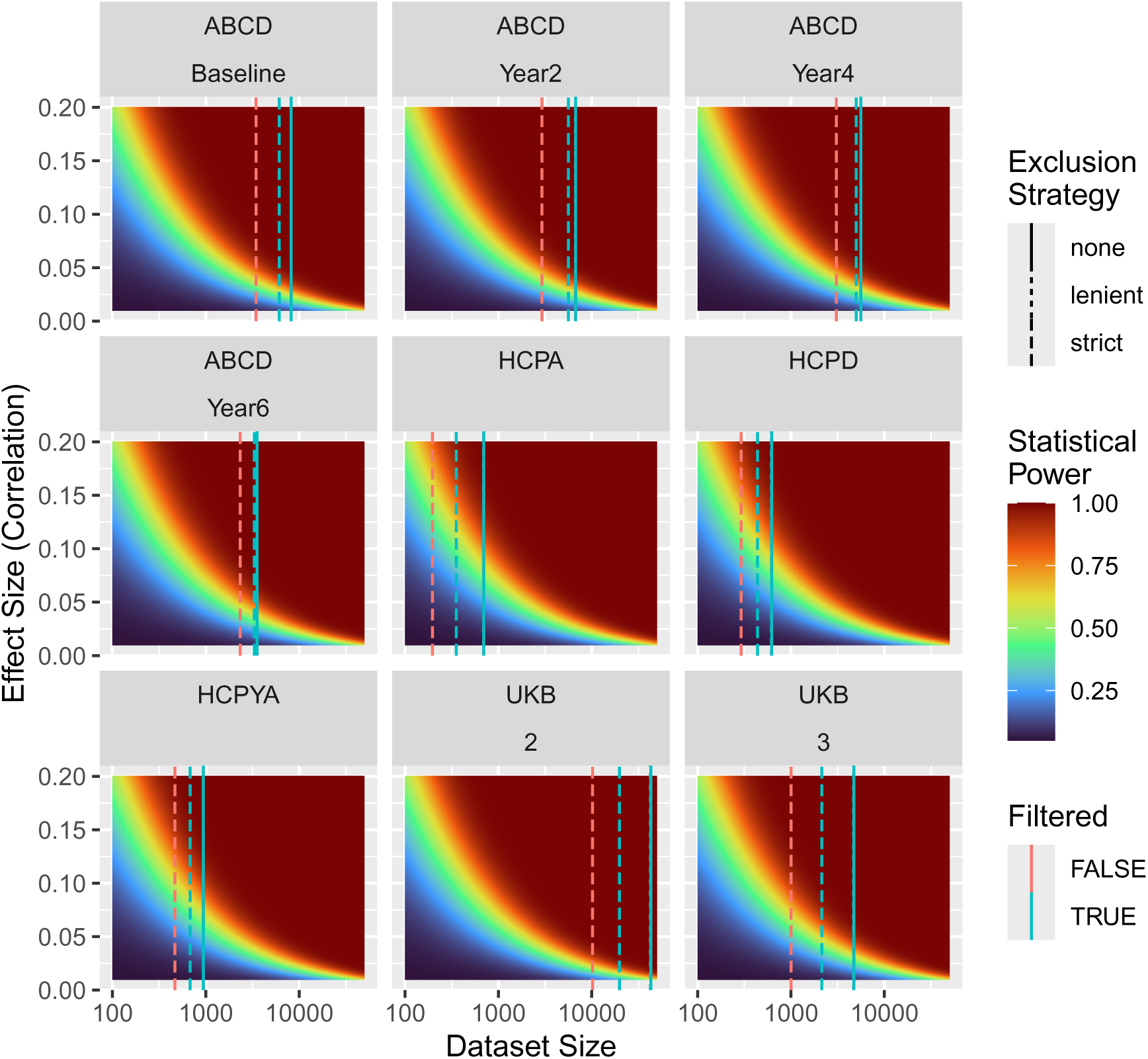
Statistical Power for Brain-Wide Association Study. FD Threshold determines rates of exclusion, as in Figure 11 and Figure 12. Note that the lenient filtering strategy never excludes many participants and so the “none” and “lenient” lines are overplotted.

To further explore the impact of exclusions on studies, we applied the economic framework presented by Ooi et al. (2025) to estimate how participant exclusion influences the cost of a study, the effect size estimated by a study, and whether the exclusion rates bias conclusions.

To estimate the cost of a study, we set a target sample size as the number of participants within each dataset. In ABCD, applying a common threshold of 0.2 mm average framewise displacement nearly doubled the required recruitment budget to reach the target usable sample size (Figure 15 a). A stricter threshold of 0.1 mm increased costs by more than an order of magnitude. In the UKB, applying the same strict threshold would result in more than a 20-fold increase in the cost required to achieve the desired sample size. Under fixed budgets, applying strict thresholds (0.1 mm) would result in less than one in four recruited participants contributing usable data (Figure 15 b). Thus, estimated budgets depend heavily on expected exclusion rates, which can vary substantially across datasets and populations.

Next, we examined the impact of exclusion rates on the framework’s estimate of predictive performance (*ρ*), using the same large-scale datasets as above. We set the *k*_*i*_ parameters to the average values derived from the ABCD resting-state data as reported by Ooi et al. (2025). This choice isolates the impact of reduced sample size under a fixed scan time, but it assumes that predictive performance does not change with the motion distribution and should therefore be viewed as a best-case scenario. In ABCD, excluding participants with average motion above 0.2 mm reduced predictive accuracy by approximately 3.93 % (Figure 15 c). Applying a stricter 0.1 mm cutoff excluded all but approximately 9.98 % of participants and reduced expected accuracy by 36.03 % of its achievable maximum. In contrast, the UK Biobank exhibited only modest declines in performance (at 0.1 mm threshold: 23.67 % of maximum accuracy), because its remaining sample size remained above the inflection point of the performance curve. When exclusions push performance below this inflection point, expected predictive performance may decline sharply even when the analytic pipeline is otherwise optimal.

Differences were observed in the relationship between training data volume and predictive performance across motion groups (Figure 17). In particular, data from the high-motion group resulted in better maximum predictive performance (binomial test for 55 features: _0.59_0.727_0.839_, *p* < 0.01), with performance measured by the correlations achieved by phenotype prediction models trained with the largest total scan time considered. This result suggests that excluding high-motion participants may alter not only sample size, but also the apparent predictability of behavioral phenotypes, presumably because the excluded participants are not behaviorally random.

**Figure 17.**
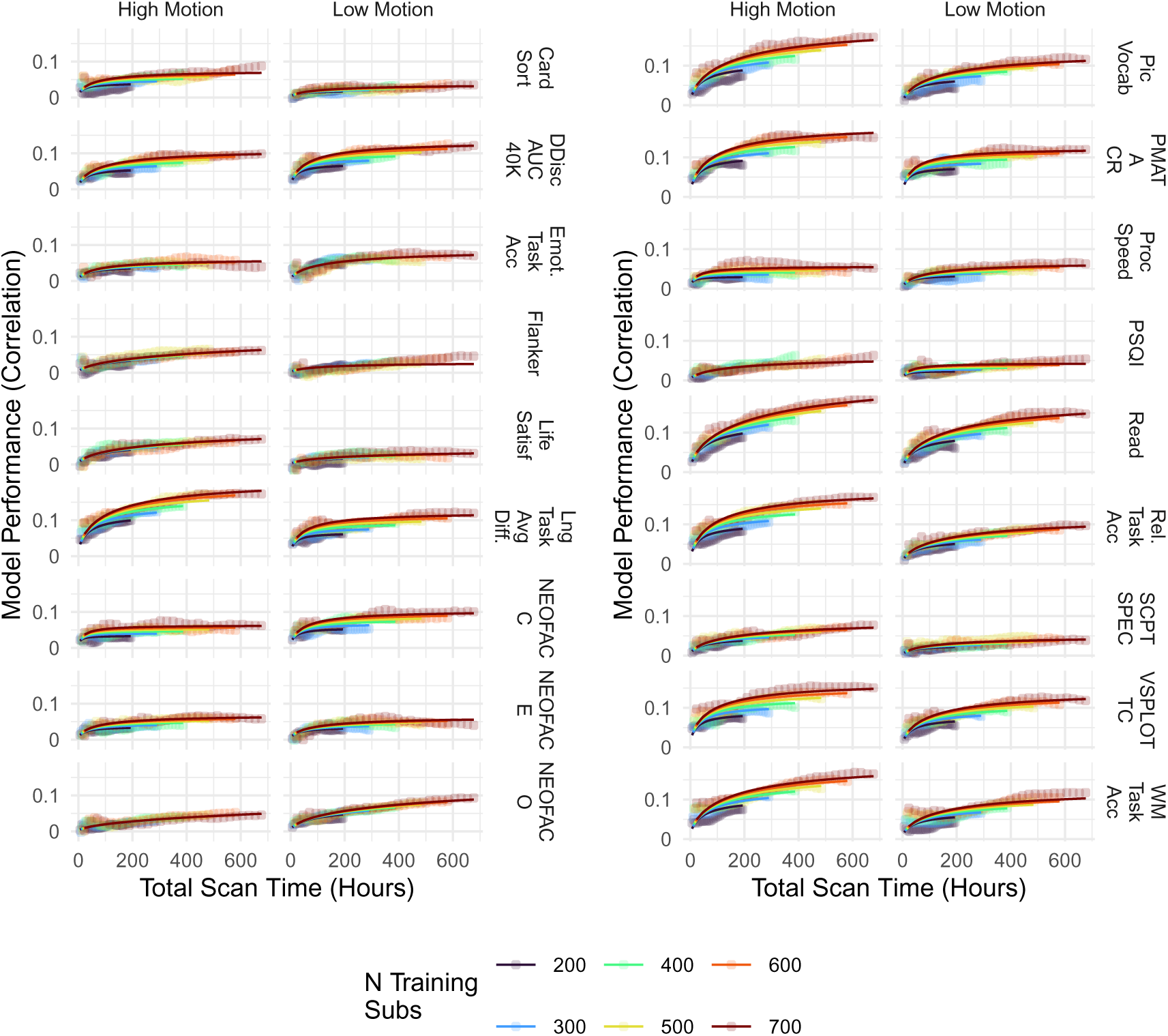
Model Performance (True and Predicted Correlation) Across Motion Groups. Using the full HCPYA dataset (MSMAll), we calculated functional connectivity as done by Ooi et al. (2025), but, to ensure sufficient time points per participant, we did not perform any censoring. Connectivity matrices were used as features in linear ridge regression models to predict phenotypes via repeated (100), nested cross-validation (restricted to the 18 phenotypes analyzed in their study). The analysis was repeated twice, once for the 800 participants with the lowest mean framewise displacement and once for the (partially overlapping) 800 participants with the highest mean framewise displacement. Points correspond to observed predictive performance, and lines depict fits of the theoretical model from Ooi et al. (2025).

This difference in asymptotic predictive performance translated into substantially different optimal designs. For example, for Fluid Intelligence (PMAT), a study that aims to achieve 80 % of maximum predictive performance with low-motion participants would be optimized with approximately 25 min scans from 3,651 individuals, whereas the optimal design with high-motion participants would require 35 min scans from 9,128 individuals. For the high-motion subgroup, the framework produces a design that requires substantially more data because performance plateaus relatively slowly in this group.

## 4 Discussion

The results of this survey reveal a sobering reality for large-scale fMRI research: standard quality-control procedures, when applied strictly, can lead to the exclusion of substantial portions of data, ultimately biasing analyses. That quality- or motion-based exclusion can negatively impact inferences by shifting sample composition is not novel (Cosgrove et al., 2022; Gard et al., 2023; Nebel et al., 2022; Peverill et al., 2025); rather, the present contribution emphasizes the scale of this problem in contemporary large datasets. For example, in the UK Biobank, strict motion thresholds could discard over 80 % of the functional imaging data. Motion was observed to be a stable trait across tasks and sessions exhibiting systematic relationships with other participant characteristics. Excluding participants based on motion therefore introduces a tension in all datasets analyzed here, a tension between biasing results by including low-quality (high motion) data and biasing results by systematically shifting the composition of sample populations.

One of the purported benefits of these datasets is their large size, both in terms of the number of participants and the depth of measurements (Falk et al., 2013). These datasets may facilitate understanding conditions that are fundamentally multimodal (Miller et al., 2016)—identifying, say, not only imaging factors that are predictive of diseases, but also clarifying how those factors can interact with fixed features (e.g., genetics), or modifiable traits with a causal role (e.g., diet). However, substantial differences between sampling and target populations present analytical challenges (Kelleher & Irani, 2026; Keyes & Westreich, 2019; *c.f.* Fry et al., 2017; Batty et al., 2020). Consider the case of the UKB and BMI, which is known to be associated with many diseases. In 2022, 64 % of UK residents were overweight or obese (Health Survey for England, 2022). Of the people with imaging, that proportion is 60 %. With a threshold of 0.2 mm, that proportion drops to 51 %. The systematic exclusion of individuals with higher BMI effectively removes variation linked to metabolic state from the analytic sample, potentially masking true associations between brain health and obesity-related physiology. This constitutes a form of collider bias (e.g., Hernán et al., 2004), a situation in which both the exposure of interest (e.g., a metabolic variable) and study inclusion are influenced by a common cause (BMI), and conditioning on inclusion occurs, as when high-BMI participants are excluded. This can introduce spurious associations or attenuate true ones, even when the original study sample was representative of the target population. Unfortunately, for neuroimaging, even original study samples are often not representative (Gard et al., 2023; Ricard et al., 2023).

We observed widespread associations with motion (and therefore rates of exclusion) between the three key demographic variables that were our focus: age, sex, BMI, and several Psychological and Life Function Variables within the HCPYA. Our observation of a relationship between motion and psychiatric variables (Figure 10) may appear to contradict an earlier report by Shi et al. (2025), who observed no relationship between motion and either externalizing or internalizing diagnoses. This discrepancy may be attributable to several sources. First, the cohorts are substantially different: Shi et al. (2025) analyzed data from the Healthy Brain Network (Alexander et al., 2017), which comprises children under the age of 16, whereas our analysis focused on young adults in the HCPYA dataset. In their dataset, the particularly strong relationship between motion and age in a developing cohort may have masked a relatively subtle association with diagnosis. Second, the coarse binarization of psychiatric state into a diagnosis of externalizing or internalizing may not be as sensitive as the continuous measurements analyzed in Figure 10.

This survey also points to a potential avenue for improving future datasets, which is to refine data collection protocols. We observed substantial differences in motion across the HCPD and ABCD datasets, although we do not know the source of these differences. Both datasets sample from pediatric cohorts, and controlling for the key characteristics considered here did not eliminate the dataset effects (age, sex, BMI, mental health factors), although additional confounding remains a possibility. One possibility is that aspects of the operating procedures contributed to the differences; although the ABCD followed several best practices for pediatric cohorts, such as the use of mock scanning and training (Casey et al., 2018; Gao et al., 2023), the ABCD scanning sessions are approximately twice as long as those in the HCPD (90 min to 120 min vs. 45 min), and, as shown here, motion tends to increase during a scanning session (see also Meissner et al., 2020). When considering the high proportions of participants that are regularly excluded from analyses of the ABCD (e.g., 40 % to 60 % Marek et al. (2019); Marek et al. (2022)), this difference between motion in the ABCD and HCPD raises the possibility that there may be substantial value gained in further improving best-practices in neuroimaging experimental operations.

For these analyses, we used the ABCD Community Collection (Feczko et al., 2021), which is released in parallel with the standard ABCD dataset. The standard ABCD dataset was observed to exhibit overall larger motion, exceeding 0.5 mm in approximately 300 scans (Section A.1). This amounts to less than 1 % of the ABCD scans analyzed here, and so we suspect that the trends shown here will be largely consistent across derivative collections.

Beyond the overall magnitude of motion, the temporal structure of motion within scans has practical consequences that deserve attention. Across datasets, motion tended to increase throughout sessions. A drift pattern means that exclusion based on average framewise displacement will disproportionately penalize longer scans, since later frames contribute disproportionately to the mean. This leads to a situation in which studies that collect more data per participant may face higher exclusion rates, thereby counteracting the statistical benefits of longer acquisition (Ooi et al., 2025). Similarly, the task-correlated motion observed across several datasets, visible as sharp spikes at button presses and elevated motion during specific task blocks, implies that exclusion rates will differ systematically between tasks even within the same participant and session. A participant retained for resting-state analysis may be excluded from task analysis, or vice versa, introducing inconsistencies in sample composition that are rarely reported. Note also that our displays of task-correlated motion, Figure 2 through Figure 7, show a conservative view of the effect, as no effort was made to account for variable task timing by aligning the motion timecourses across participants. These patterns collectively argue for exclusion criteria adapted to scan type, task structure, and acquisition length, rather than applied uniformly.

Although there is substantial motion in these datasets, and although commonly applied thresholds entail exceptionally high rates of participant exclusion, this does not necessarily mean most of these datasets are unusable. This qualification applies most strongly to adult cohorts, where respiratory pseudo-motion is a major contributor to framewise displacement. In all datasets, we observed a distinct spectral peak in the motion parameters at respiratory frequencies (approximately 0.3 Hz), particularly in the phase-encoding direction. This is consistent with “pseudo-” or “apparent” motion—an artifact caused not by the head moving, but by the chest wall expanding and altering the B0 magnetic field.^1^ Motion that is only apparent may have a less substantial impact on data quality as compared with mechanical motion (Fair et al., 2020; Phạm et al., 2023). Consistent with earlier work (Phạm et al., 2023), a data-driven method of censoring (DVARS) had a larger effect on functional connectivity estimates than censoring based on framewise displacement, suggesting that not all high-motion frames necessarily corrupt the timeseries (e.g., when motion is only apparent), and that some high-motion participants may still contribute usable data.

Overall, the relative risk of bias from including versus excluding high-motion participants remains unresolved. Consider the difference in predictive performance between high- versus low-motion participants; high-motion participants tended to yield higher predictive performance (Figure 17). This pattern is not surprising, as head motion is not purely random noise but is systematically associated with behavioral phenotypes, providing learnable structure that prediction models can exploit. For example, average motion has a rank correlation of −0.17 with Card Sorting performance (Card Sort) and −0.12 with the area under the delay discounting curve for evaluation of $40k (DDisc AUC 40k). Importantly, this result does not indicate superior data quality in high-motion participants. Rather, it is consistent with exclusion of high-motion individuals removing a non-random, behaviorally distinct subgroup.

Finally, it is important to contextualize the high exclusion rates reported here by noting that strict FD thresholds do not account for the efficacy of modern denoising techniques. For example, the UK Biobank pipeline utilizes ICA-FIX, which automatically identifies and removes noise components from the data (Griffanti et al., 2014). This removal allows ICA-FIX to recover usable signal from scans that might otherwise be discarded based solely on raw motion metrics, performing a sort of “soft-censoring” on the data. This suggests that although 80 % of UKB participants might fail a “strict” motion criterion, a substantially larger fraction may still contribute valid data after aggressive denoising. Indeed, some evidence suggests that even high-motion frames substantially improve functional connectivity estimates (Mejia et al., 2026). At present, however, the usable limit of data heavily influenced by motion remains difficult to quantify. Consider also that the HCPYA dataset, one of the field’s most widely used resources, was recently re-released with a new denoising procedure that augments ICA-FIX with a temporal ICA-based approach (Glasser et al., 2018; Yang et al., 2024), underscoring that ICA-FIX alone does not remove all sources of structured noise. Additional research is needed to establish how much motion can be tolerated under contemporary preprocessing pipelines and how such denoising affects downstream analyses.

### 4.1 Recommendations

The findings presented here point towards several concrete changes in how the field approaches motion-related quality control. We outline several recommendations to improve both data quality and the representativeness of neuroimaging samples.

#### 4.1.1 Report demographics of participants excluded due to motion

Studies must explicitly report the demographics of participants excluded due to motion, and not merely the exclusion rate. “Data quality” is not a neutral filter; it is a demographic sieve whose consequences are rarely made visible. For guidelines on which demographic variables to include, we defer to the Human Brain Mapping Committee on Best Practices in Data Analysis and Sharing report (Nichols et al., 2016). Pre-registering quality-control criteria, including the motion threshold and the demographic variables to be reported, would further reduce researchers’ degrees of freedom and improve the reproducibility of downstream findings.

#### 4.1.2 With modern scanning sequences, prefer data-driven methods of exclusion over strict motion filters

In modern datasets with sub-second TRs, a substantial proportion of apparent motion reflects respiratory pseudo-motion rather than mechanical head movement, particularly in adult cohorts.

Applying fixed framewise displacement thresholds under these conditions can lead to unnecessary exclusion. Where motion parameters are used to identify frames or participants for removal, data-driven methods based on signal properties are preferable to fixed FD cutoffs (e.g., DVARS-based censoring, Phạm et al., 2023). If data-driven methods are not available, applying a respiratory notch filter to motion parameters before thresholding can recover adult participants lost to pseudo-motion. This strategy should be used cautiously in pediatric cohorts, where motion is predominantly mechanical and respiratory filtering may obscure genuine data quality issues.

#### 4.1.3 Clarify target populations and evaluate reweighting strategies

Motion-corrupted data biases results, and yet motion-based exclusion disproportionately removes individuals who are already underrepresented in neuroimaging (Dhamala et al., 2025; Ricard et al., 2023). Where sample sizes permit, inverse probability weighting or related post-stratification techniques can be used to improve the external validity of analyses in the presence of structured exclusion (Gard et al., 2023; LeWinn et al., 2017). It is important that the feasibility of such approaches should be assessed at the study design stage, and not applied as an afterthought.

### 4.2 Limitations

Several limitations of the present analysis should be acknowledged. First, framewise displacement was used as the primary motion metric throughout. While FD is the most widely used measure and facilitates comparisons across studies, it is not the only available metric, and its relationship to data quality is imperfect, particularly at sub-second TRs. Second, the cost and predictive performance estimates relied on an extrapolation with the framework by Ooi et al. (2025) that extended well beyond the amount of scan time on which the underlying model was fit, and so those estimates should be interpreted with caution. Third, motion is related to multidimensional profiles (Bolton et al., 2020), which may color the univariate relationships between motion and participant characteristics reported here. Finally, the present survey is descriptive rather than prescriptive; we quantify the consequences of commonly used motion thresholds and join others in recommending replacing them with data-driven criteria, but we do not derive optimal thresholds, which will necessarily depend on the scientific question, the population, and the preprocessing pipeline employed.

### 4.3 Conclusions

Motion in fMRI is often treated as a technical nuisance—noise to be modeled, flagged, and removed before analysis. The results of this survey suggest that this framing is incomplete. Across six large, publicly available datasets spanning nearly the entire human lifespan, motion-based exclusion is both substantial and demographically structured. Under commonly used strict thresholds, large fractions of participants are removed, and these removals are not random.

Younger and older individuals, those with higher BMI, and those with motion-associated conditions are disproportionately excluded, reshaping the analytic sample before analysis begins. This matters because the value of these datasets lies in their scale and diversity. Motion-based exclusion undermines both. A threshold that discards most of a dataset, or systematically underrepresents specific groups, does not simply reduce sample size; it alters the population under study, often without explicit acknowledgment. At the same time, this is not an argument against quality control. Motion can meaningfully degrade fMRI data, and mitigating its effects remains essential. The challenge is to improve data quality without treating representativeness as expendable. Approaches such as respiratory filtering, data-driven exclusion criteria, and prospective motion mitigation offer practical ways forward. Equally important is transparency, and reporting the demographic composition of excluded participants should be standard practice. As neuroimaging moves toward population-scale science, quality-control decisions will increasingly shape what the field can observe. Motion thresholds are not neutral, and treating them as such risks embedding systematic biases into the literature that larger sample sizes alone cannot resolve.

## Data and Code Availability

Analyses relied on open data from the Human Connectome Project, available for download from the HCP website humanconnectome.org.

The data that support the findings of this study are available from UK Biobank. Restrictions apply to the availability of these data, which were used under license for this study. Data are available from https://www.ukbiobank.ac.uk/enable-your-research/apply-for-access with the permission of UK Biobank.

The de-identified ABCD data and data dictionaries are available through the NIMH Data Archive (NDA) following approval of a Data Use Certification (DUC). Data releases are hosted on the NIH Brain Development Cohorts.

The SpaceTop motion data is available on OpenNeuro as ds007070.

Code to reproduce analyses is available on GitHub: psadil/motion_fmri_survey.

## Author Contributions

**Conceptualization**: PS, BA, ACM, ASC, JC, FVF, LEI, MAJ, AK, MBN, JJP, HIS, JS, AMS, MAL; **Data Curation**: PS, BA, ACM, LEI, AK, JS; **Formal Analysis**: PS, BA, ACM, JC, LEI, AK, JS; **Funding Acquisition**: MAL; **Project Administration**: PS, MAL; **Resources**: PS; **Software**: PS, BA, ACM, LEI, AK, JS; **Supervision**: MAL; **Visualization**: PS, BA, ACM, LEI, AK, MBN, JS, TDW, MAL; **Writing – Original Draft**: PS, BA, LEI, JS; **Writing – Review & Editing**: PS, ACM, ASC, MBN, JJP, JS, TDW, MAL.

## Acknowledgements

Data were provided by the Human Connectome Project, WU-Minn Consortium (Principal Investigators: David Van Essen and Kamil Ugurbil; 1U54MH091657), funded by the 16 NIH Institutes and Centers that support the NIH Blueprint for Neuroscience Research, and by the McDonnell Center for Systems Neuroscience at Washington University.

Research reported in this publication was supported by the National Institute On Aging of the National Institutes of Health under Award Number U01AG052564. The content is solely the responsibility of the authors and does not necessarily represent the official views of the National Institutes of Health.

Research reported in this publication was supported by the National Institute Of Mental Health of the National Institutes of Health under Award Number U01MH109589. The content is solely the responsibility of the authors and does not necessarily represent the official views of the National Institutes of Health.

This research has been conducted using data from UK Biobank, a major biomedical database (Project ID: 33278). We are grateful to UK Biobank and the UK Biobank participants for making the resource data possible, and to the data processing team at Oxford University for sharing the processed data. The UK Biobank imaging project is funded by the Medical Research Council and the Wellcome Trust.

Data used in the preparation of this article were obtained from the Adolescent Brain Cognitive DevelopmentSM (ABCD) Study (https://abcdstudy.org), held in the NIMH Data Archive (NDA). This is a multisite, longitudinal study designed to recruit more than 10,000 children age 9-10 and follow them over 10 years into early adulthood. The ABCD Study® is supported by the National Institutes of Health and additional federal partners under award numbers U01DA041048, U01DA050989, U01DA051016, U01DA041022, U01DA051018, U01DA051037, U01DA050987, U01DA041174, U01DA041106, U01DA041117, U01DA041028, U01DA041134, U01DA050988, U01DA051039, U01DA041156, U01DA041025, U01DA041120, U01DA051038, U01DA041148, U01DA041093, U01DA041089, U24DA041123, U24DA041147.

A full list of supporters is available at https://abcdstudy.org/federal-partners.html. A listing of participating sites and a complete listing of the study investigators can be found at https://abcdstudy.org/consortium_members/. ABCD consortium investigators designed and implemented the study and/or provided data but did not necessarily participate in the analysis or writing of this report. This manuscript reflects the views of the authors and may not reflect the opinions or views of the NIH or ABCD consortium investigators.

## Supplementary Materials

### A.1 Motion in ABCD vs. ABCC

**Figure A1.**
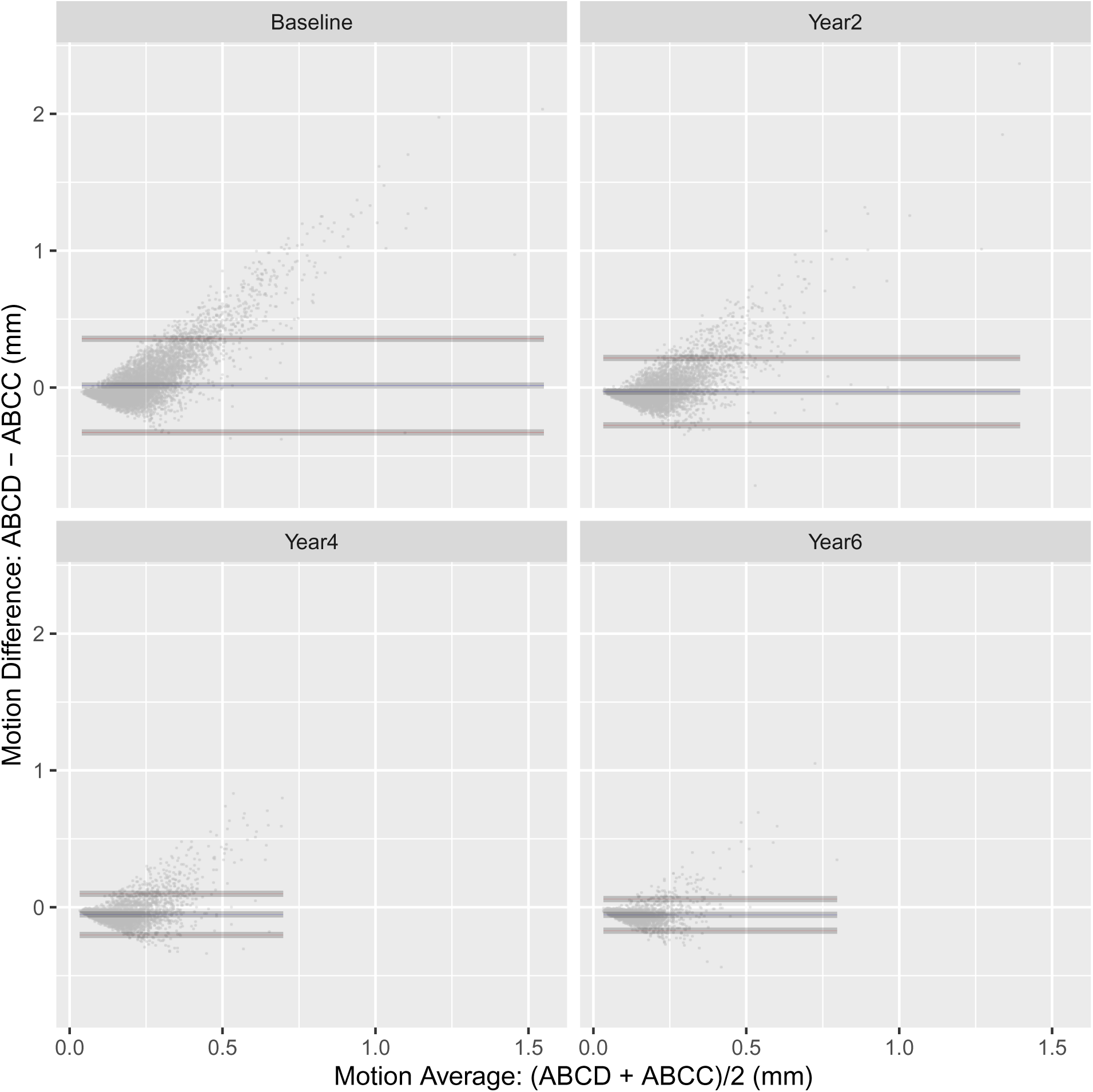
Comparison of Motion in Resting-State Scans Across ABCC and ABCD. Each point corresponds to a participant. The blue ribbons mark average differences, and the red ribbons are the limits of agreement (approximately 2 standard deviations of the difference). Ribbon widths indicate 95 % confidence intervals for the estimates. Motion in the ABCD scans was pulled from the `mr_y_qc mot rsfmri mot_mean` variable (average framewise displacement in resting-state fMRI).

### A.2 Motion Spectral Frequencies

In all plots below, only the first run of a task is included per session. Spectra are calculated as in Fair et al. (2020). Briefly, power spectral densities, *p*, were estimated after detrending with an adaptive, multitaper filter (NFFT=512, NW=8), scaled by 10 log(*p*), z-scored, linearized, converted to percentages, and truncated at 0.1 % and 96 %. For a detailed description of the procedure see Fair et al. (2020). Within each panel, rows correspond to participants ordered by average framewise displacement. Because a panel is far too short to resolve individual participants, spectra from datasets with more than 256 participants were averaged within 256 equally sized bins of that ordering; this discards only vertical resolution that could not have been displayed at print resolution. SpaceTop has fewer than 256 participants and so is shown unbinned, with one row per participant.

**Figure A2.**
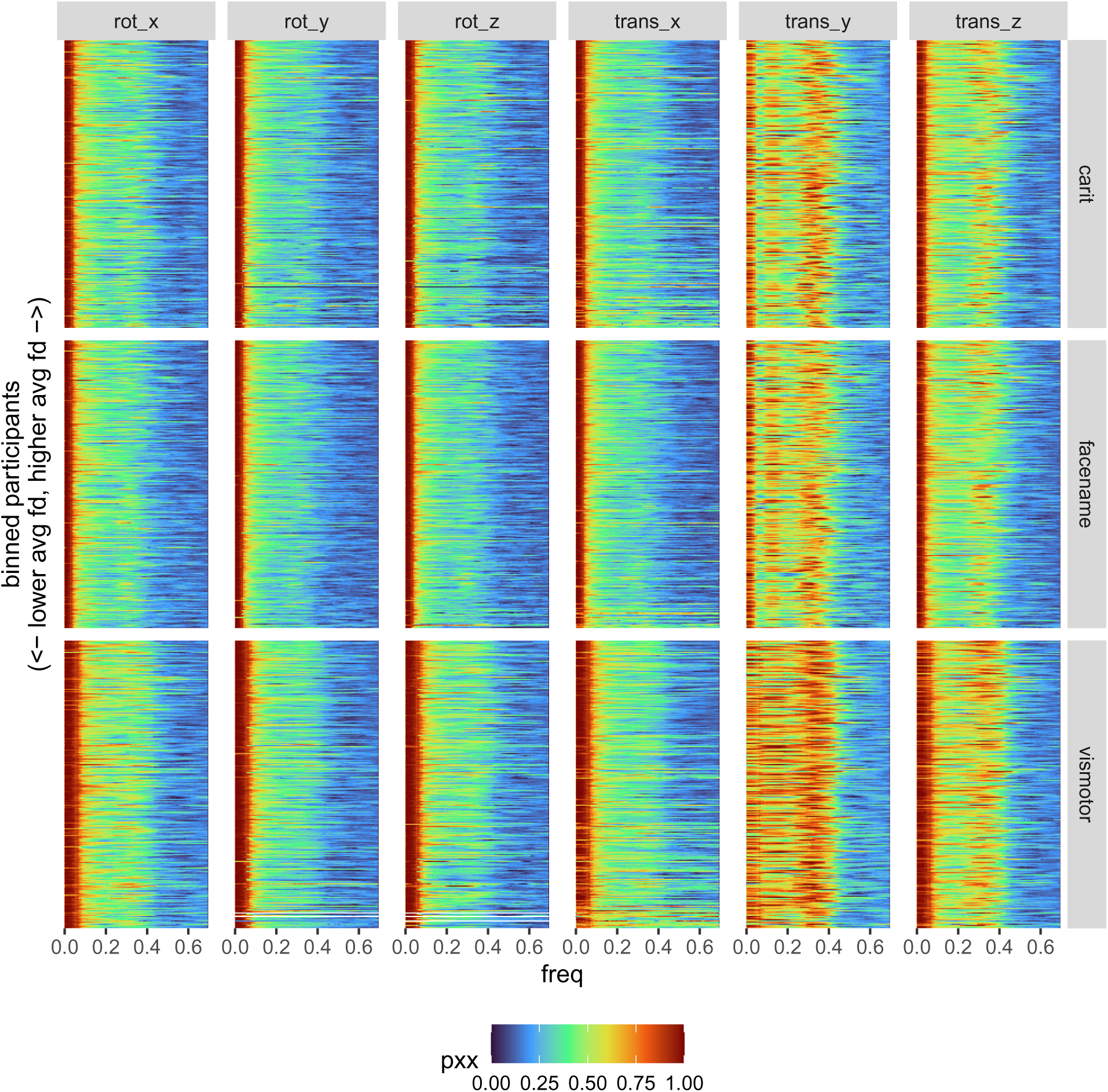
Motion Spectrum in HCPA.

**Figure A3.**
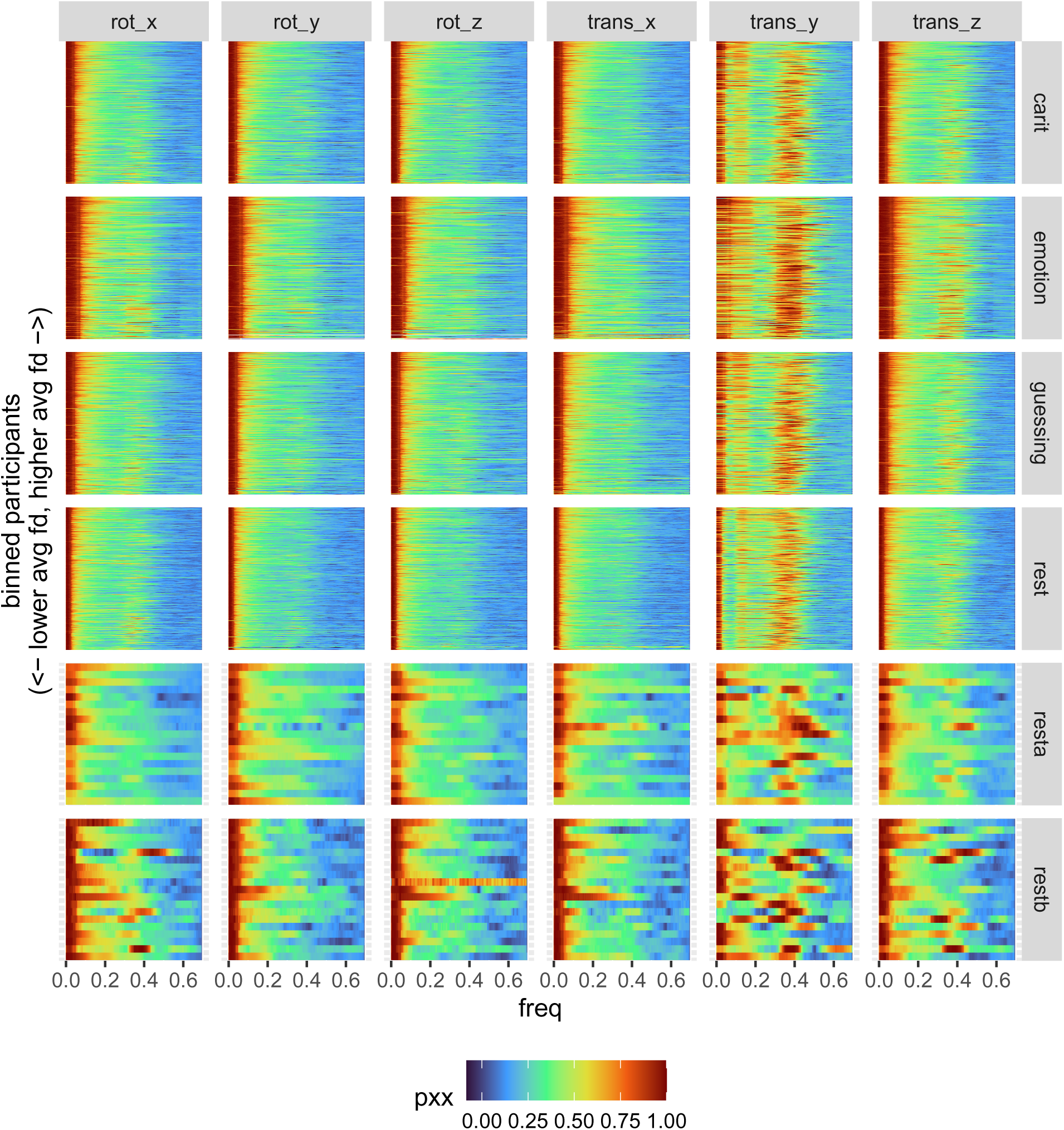
Motion Spectrum in HCPD.

**Figure A4.**
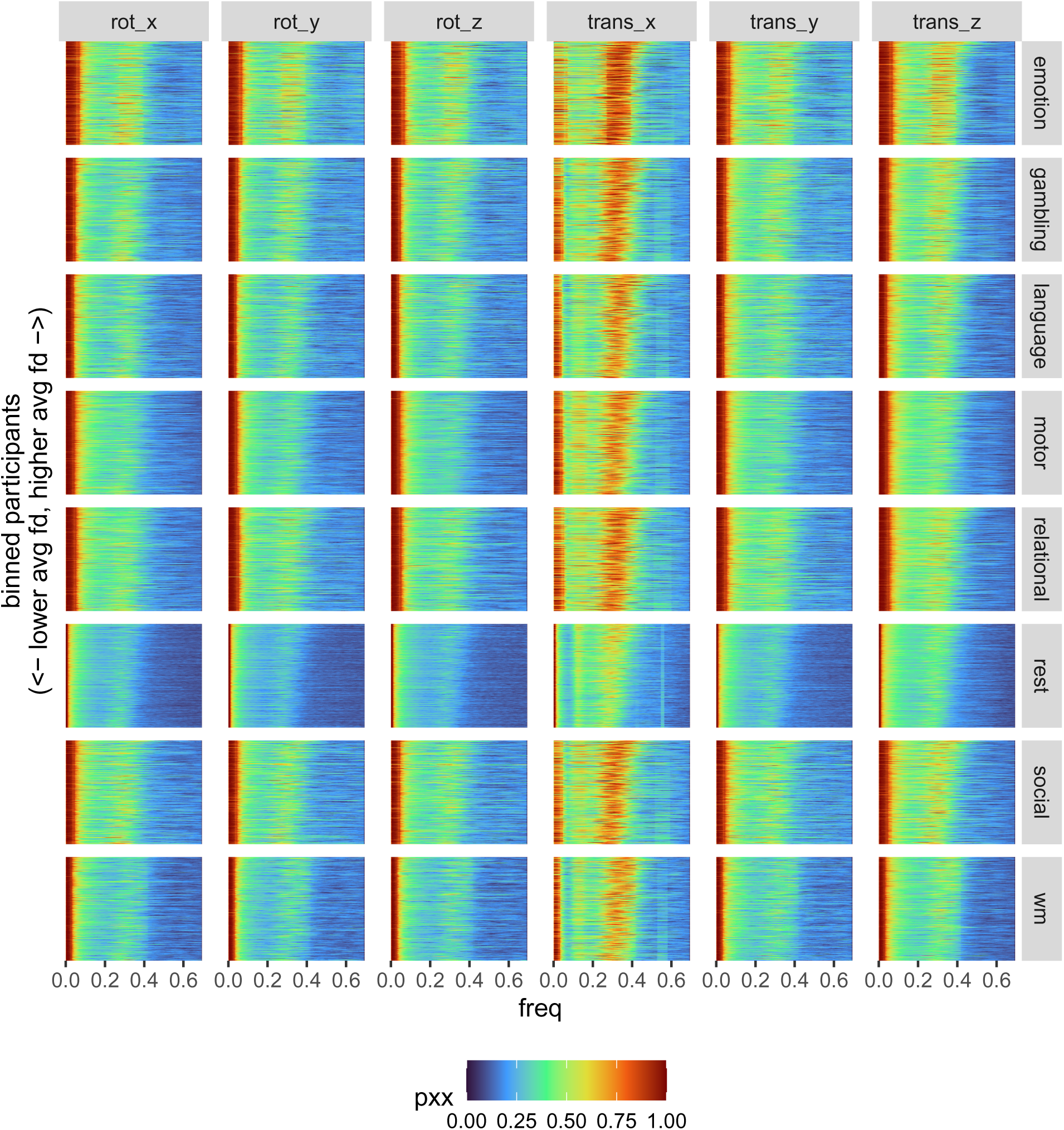
Motion Spectrum in HCPYA.

**Figure A5.**
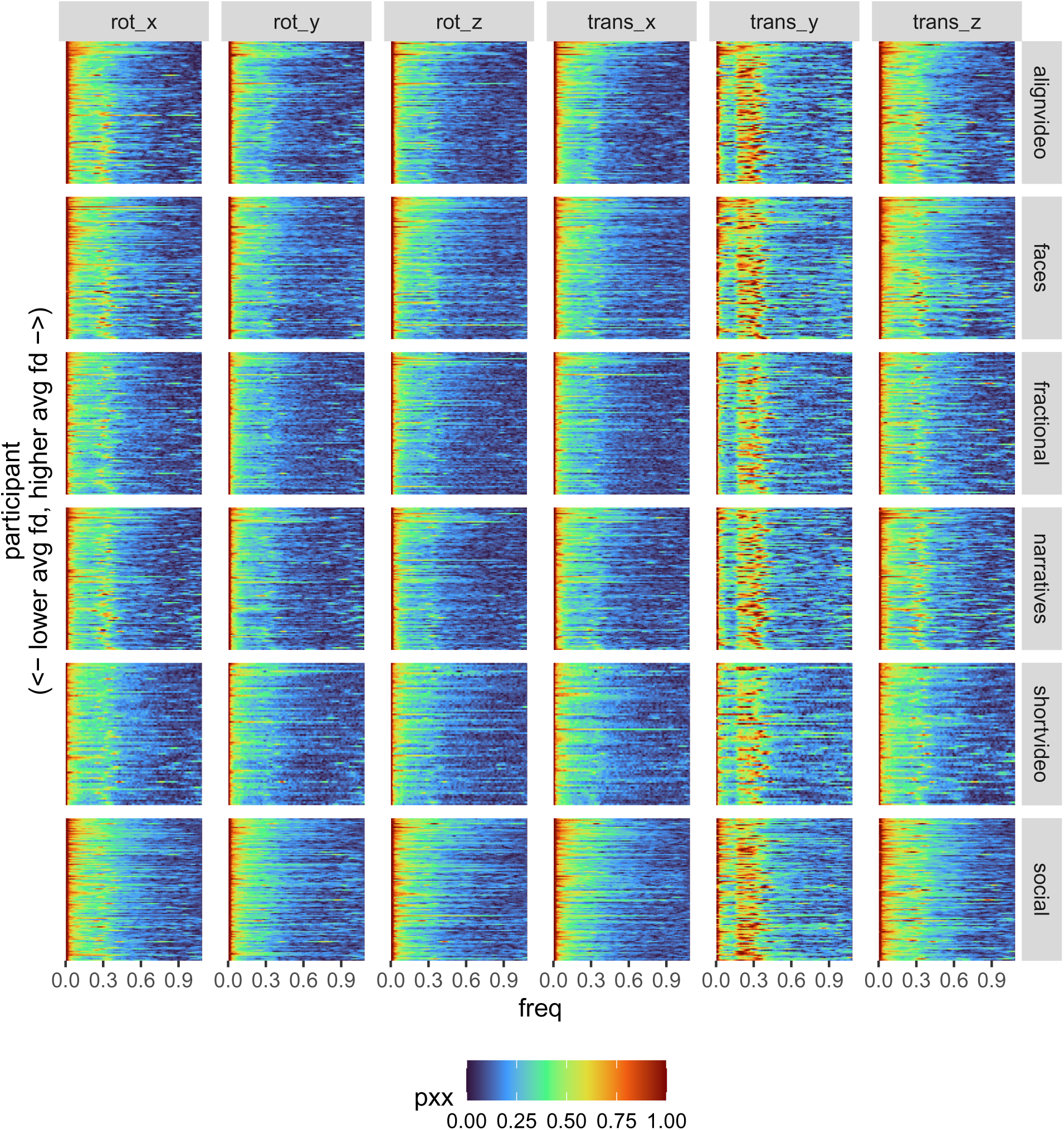
Motion Spectrum in SpaceTop.

**Figure A6.**
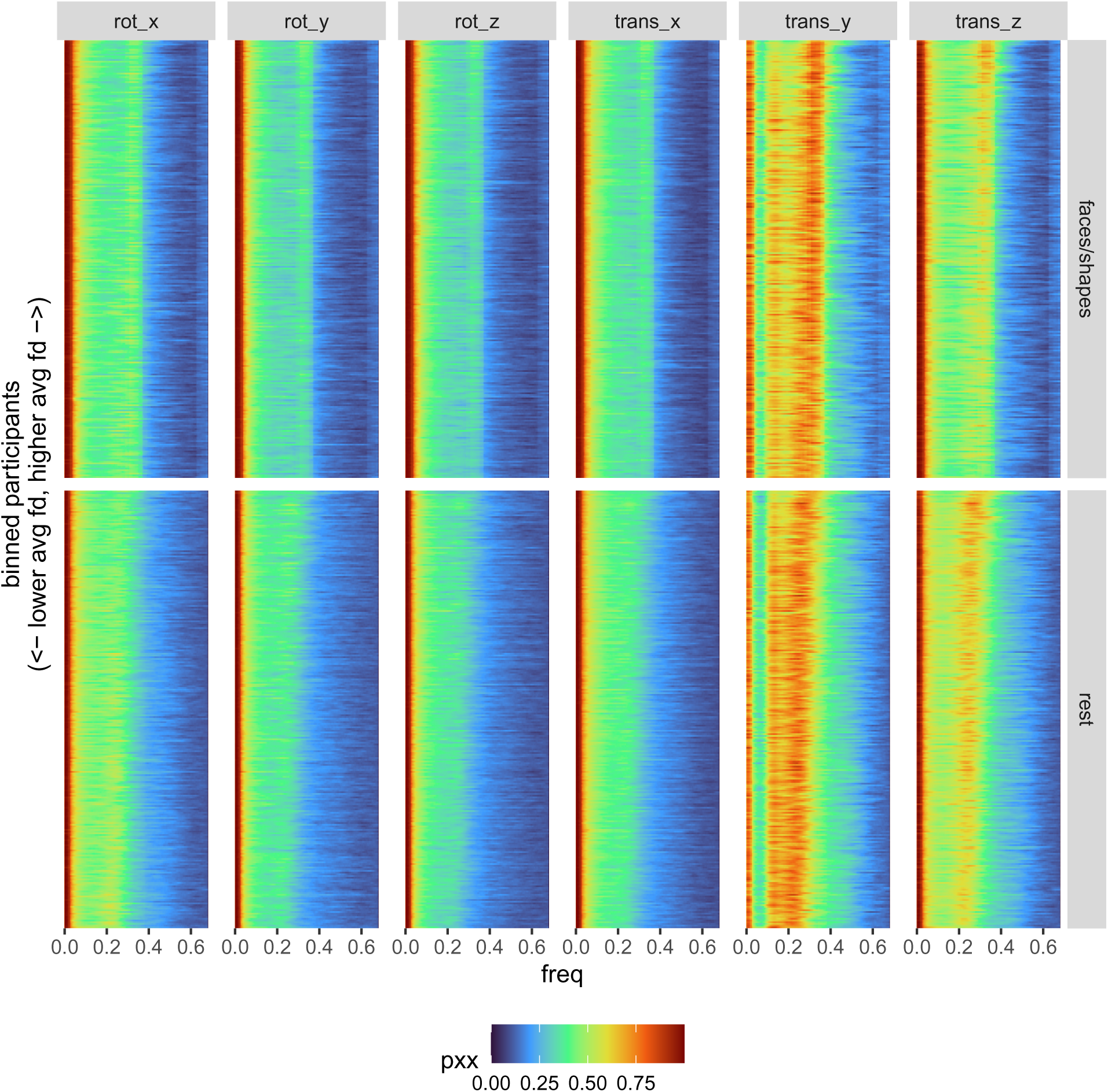
Motion Spectrum in UKB (Repeat Session Only).

**Figure A7.**
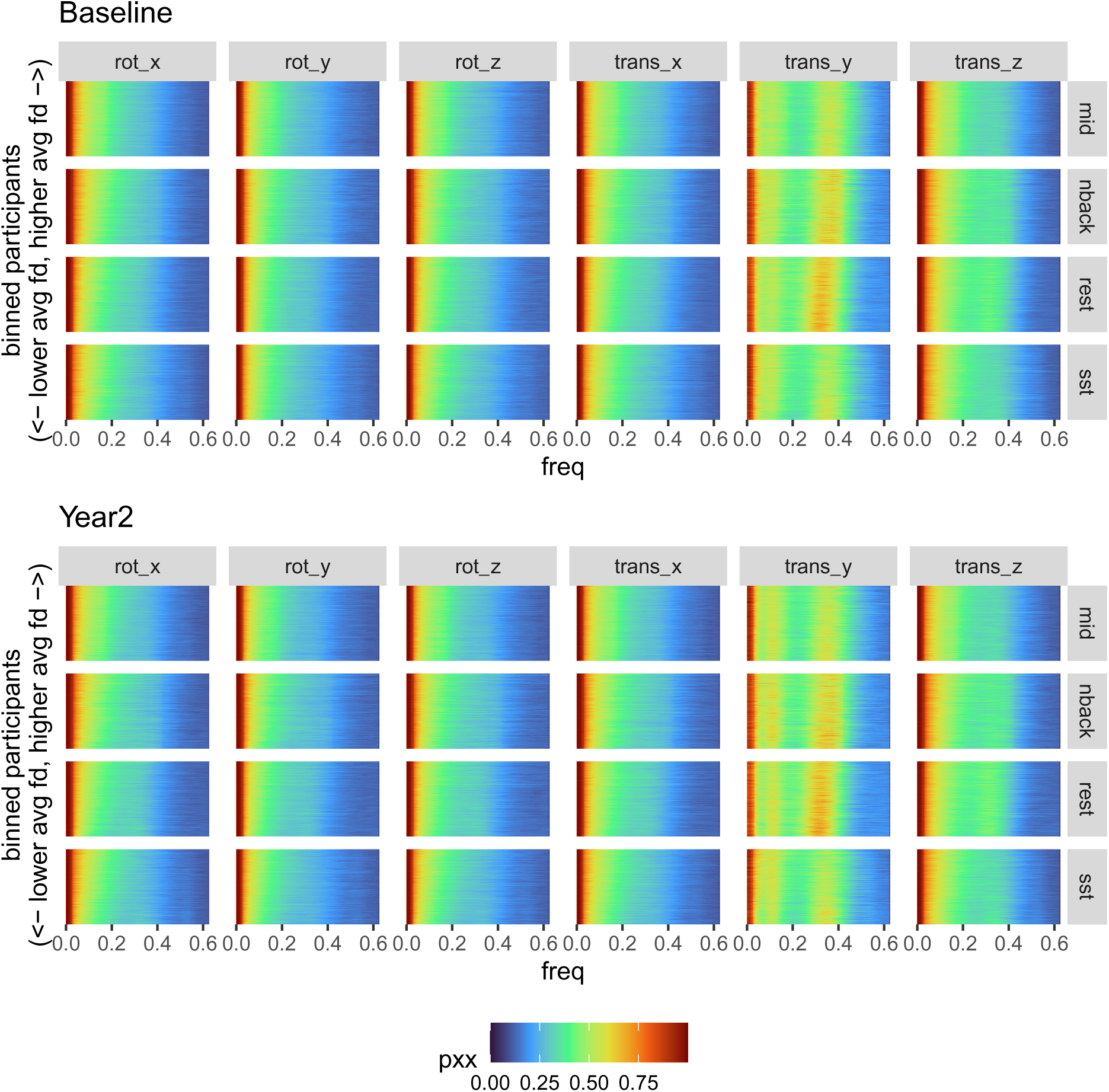
Motion Spectrum in ABCD (First Two Sessions).

**Figure A8.**
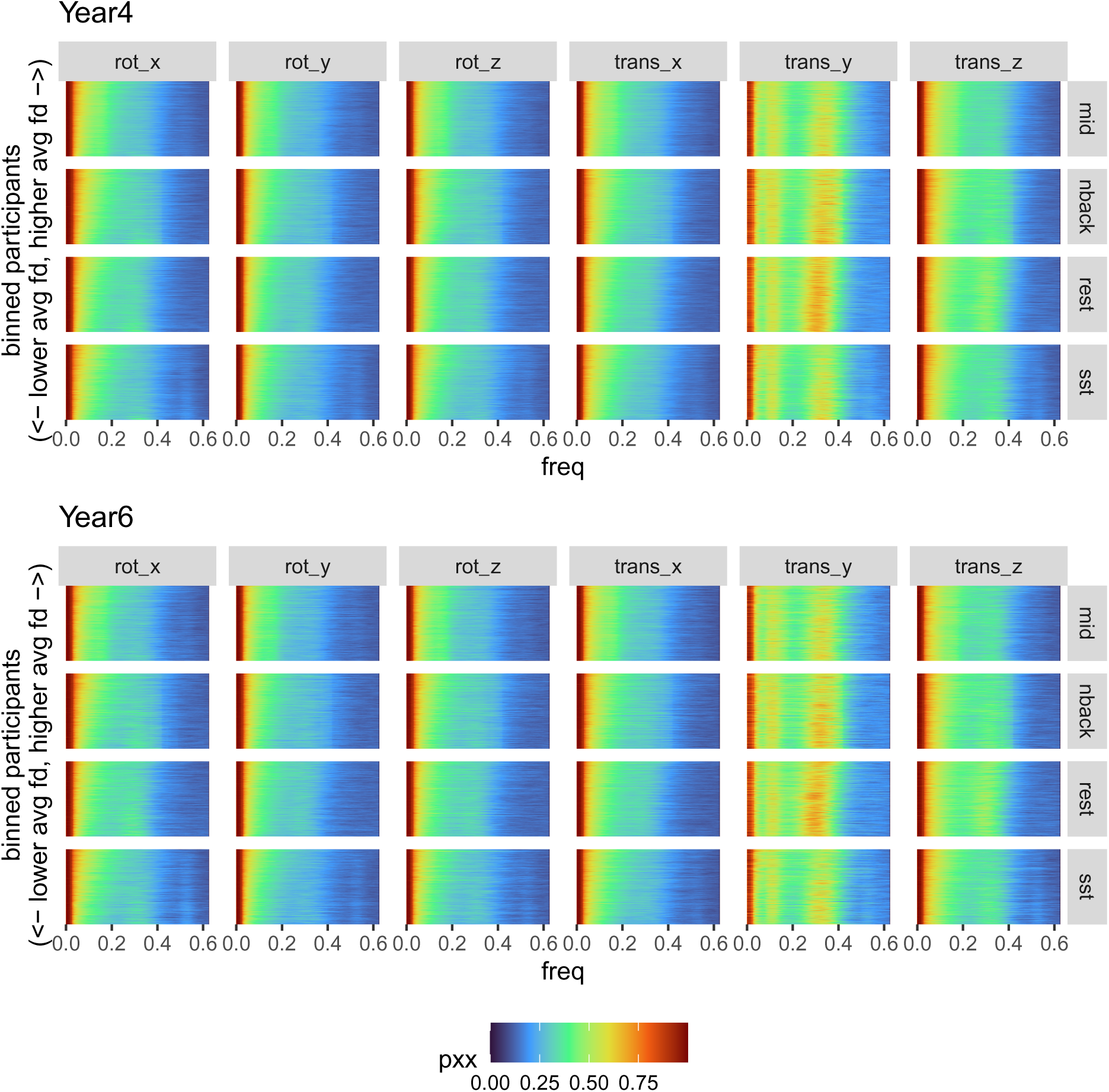
Motion Spectrum in ABCD (Final Two Sessions).

### A.3 Model Fit

**Table A1.**
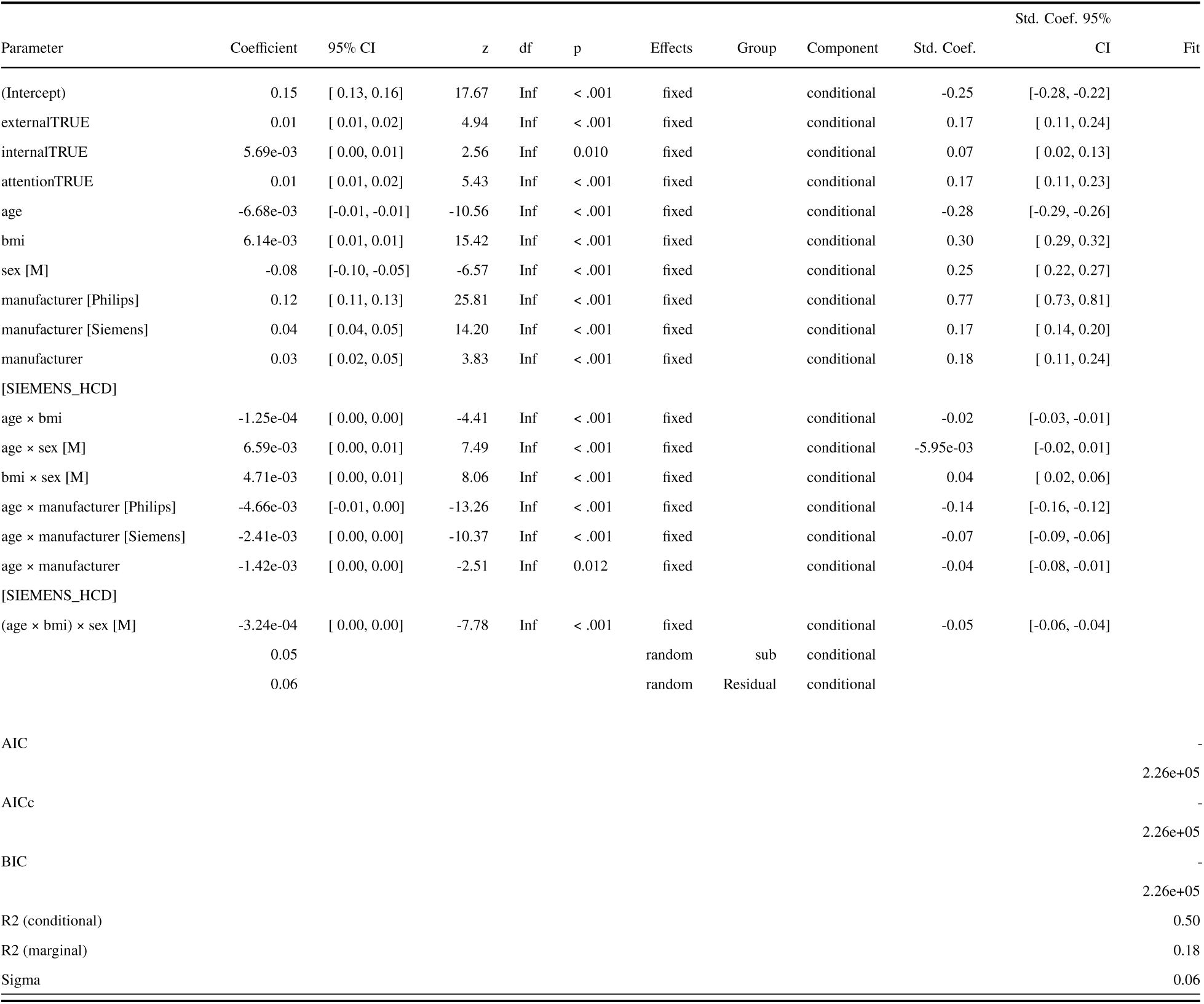
Linear Mixed Effects Model Summary of Average Motion in Resting-State Scans for Comparison of HCPD and ABCD. The model formula was `fd ∼ external + internal + attention + age * bmi * sex + manufacturer + manufacturer:age + (1 | sub)`, where `fd` is runwise average framewise displacement (in ABCD and HCPD only), `external`, `internal`, and `attention` are binary factors indicating symptoms, and `manufacturer` is a factor indicating the manufacturer of the scanner used for scanning. Manufacturer was included to account for differences in the acquisition parameters (Hagler Jr et al., 2019). All HCPD scans were collected on a Siemens scanner, and so, for convenience of constructing a contrast for dataset, the manufacturer in the HCPD dataset was set to Siemens-HCD. The model was fit using `glmmTMB` (McGillycuddy et al., 2025).

**Table A2.**
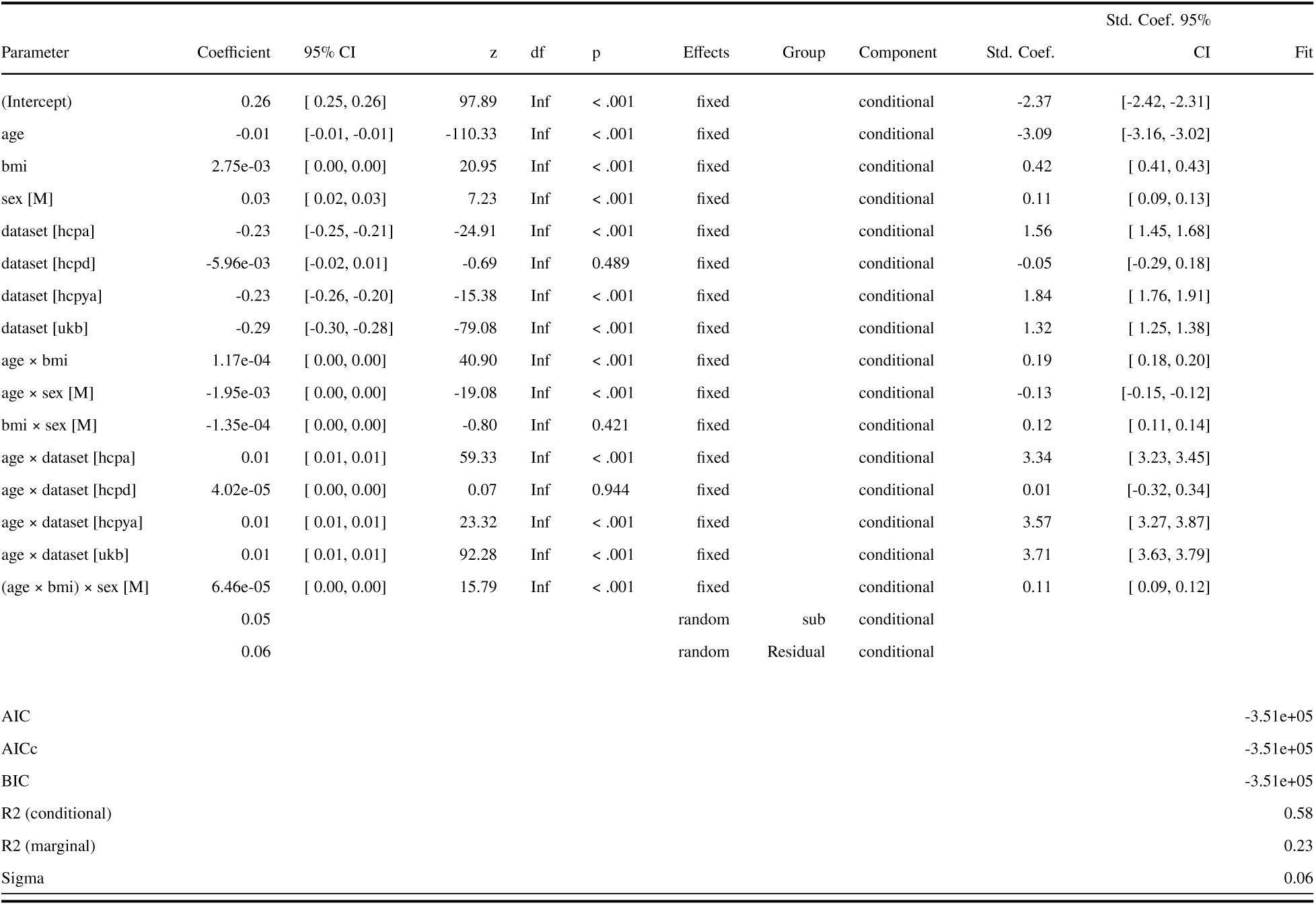
Linear Mixed Effects Model Summary of Average Motion in Resting-State Scans. The model formula was `fd ∼ age * bmi * sex + dataset + dataset:age + (1 | sub)`, where `fd` is runwise average framewise displacement (all datasets included). The model was fit using `glmmTMB` (McGillycuddy et al., 2025).

## Footnotes

1 In contrast to the report by Power et al. (2019), we observed this phenomenon in datasets that included children (ABCD, HCPD, see also Fair et al., 2020). It has been hypothesized that children may exhibit less pseudo-motion due to their smaller chest cavities. However, pseudo-motion is harder to identify in the presence of real motion, which is typically high in children. That is, we speculate that Power et al. (2019) may have observed minimal pseudo-motion in their younger cohort because of that cohort’s propensity to move frequently, and their relatively small sample size (<100) did not include enough low-motion children to uncover pseudo-motion.

## References

Achenbach, T. M. (2009). The achenbach system of empirically based assessment (ASEBA): Development, findings, theory, and applications. University of Vermont, Research Center for Children, Youth, & Families.

Afyouni, S., & Nichols, T. E. (2018). Insight and inference for DVARS. NeuroImage, 172, 291–312. 10.1016/j.neuroimage.2017.12.098

Alexander, L. M., Escalera, J., Ai, L., Andreotti, C., Febre, K., Mangone, A., Vega-Potler, N., Langer, N., Alexander, A., Kovacs, M., et al. (2017). An open resource for transdiagnostic research in pediatric mental health and learning disorders. Scientific Data, 4(1), 170181. 10.1038/sdata.2017.181

Alfaro-Almagro, F., Jenkinson, M., Bangerter, N. K., Andersson, J. L., Griffanti, L., Douaud, G., Sotiropoulos, S. N., Jbabdi, S., Hernandez-Fernandez, M., Vallee, E., et al. (2018). Image processing and quality control for the first 10,000 brain imaging datasets from UK biobank. NeuroImage, 166, 400–424.

Barch, D. M., Albaugh, M. D., Avenevoli, S., Chang, L., Clark, D. B., Glantz, M. D., Hudziak, J. J., Jernigan, T. L., Tapert, S. F., Yurgelun-Todd, D., Alia-Klein, N., Potter, A. S., Paulus, M. P., Prouty, D., Zucker, R. A., & Sher, K. J. (2018). Demographic, physical and mental health assessments in the adolescent brain and cognitive development study: Rationale and description. Developmental Cognitive Neuroscience, 32, 55–66. 10.1016/j.dcn.2017.10.010

Barch, D. M., Burgess, G. C., Harms, M. P., Petersen, S. E., Schlaggar, B. L., Corbetta, M., Glasser, M. F., Curtiss, S., Dixit, S., Feldt, C., et al. (2013). Function in the human connectome: Task-fMRI and individual differences in behavior. Neuroimage, 80, 169–189. 10.1016/j.neuroimage.2013.05.033

Batty, G. D., Gale, C. R., Kivimäki, M., Deary, I. J., & Bell, S. (2020). Comparison of risk factor associations in UK Biobank against representative, general population based studies with conventional response rates: Prospective cohort study and individual participant meta-analysis. BMJ, m131. 10.1136/bmj.m131

Bolton, T. A. W., Kebets, V., Glerean, E., Zöller, D., Li, J., Yeo, B. T. T., Caballero-Gaudes, C., & Van De Ville, D. (2020). Agito ergo sum: Correlates of spatio-temporal motion characteristics during fMRI. NeuroImage, 209, 116433. 10.1016/j.neuroimage.2019.116433

Bookheimer, S. Y., Salat, D. H., Terpstra, M., Ances, B. M., Barch, D. M., Buckner, R. L., Burgess, G. C., Curtiss, S. W., Diaz-Santos, M., Elam, J. S., et al. (2019). The lifespan human connectome project in aging: An overview. Neuroimage, 185, 335–348. 10.1016/j.neuroimage.2018.10.009

Casey, B. J., Cannonier, T., Conley, M. I., Cohen, A. O., Barch, D. M., Heitzeg, M. M., Soules, M. E., Teslovich, T., Dellarco, D. V., Garavan, H., Orr, C. A., Wager, T. D., Banich, M. T., Speer, N. K., Sutherland, M. T., Riedel, M. C., Dick, A. S., Bjork, J. M., Thomas, K. M., … Dale, A. M. (2018). The Adolescent Brain Cognitive Development (ABCD) study: Imaging acquisition across 21 sites. Developmental Cognitive Neuroscience, 32, 43–54. 10.1016/j.dcn.2018.03.001

Chyzhyk, D., Varoquaux, G., Milham, M., & Thirion, B. (2022). How to remove or control confounds in predictive models, with applications to brain biomarkers. GigaScience, 11, giac014. 10.1093/gigascience/giac014

Cicchetti, D. V., & Sparrow, S. A. (1981). Developing criteria for establishing interrater reliability of specific items: applications to assessment of adaptive behavior. American Journal of Mental Deficiency, 86(2), 127–137.

Ciric, R., Wolf, D. H., Power, J. D., Roalf, D. R., Baum, G. L., Ruparel, K., Shinohara, R. T., Elliott, M. A., Eickhoff, S. B., Davatzikos, C., Gur, R. C., Gur, R. E., Bassett, D. S., & Satterthwaite, T. D. (2017). Benchmarking of participant-level confound regression strategies for the control of motion artifact in studies of functional connectivity. NeuroImage, 154, 174–187. 10.1016/j.neuroimage.2017.03.020

Compton, W. M., Dowling, G. J., & Garavan, H. (2019). Ensuring the best use of data: The adolescent brain cognitive development study. JAMA Pediatrics, 173(9), 809–810. 10.1001/jamapediatrics.2019.2081

Cosgrove, K. T., McDermott, T. J., White, E. J., Mosconi, M. W., Thompson, W. K., Paulus, M. P., Cardenas-Iniguez, C., & Aupperle, R. L. (2022). Limits to the generalizability of resting-state functional magnetic resonance imaging studies of youth: An examination of ABCD study® baseline data. Brain Imaging and Behavior, 16(4), 1919–1925. 10.1007/s11682-022-00665-2

Dhamala, E., Ricard, J. A., Uddin, L. Q., Galea, L. A., Jacobs, E. G., Yip, S. W., Yeo, B. T., Chakravarty, M. M., & Holmes, A. J. (2025). Considering the interconnected nature of social identities in neuroimaging research. Nature Neuroscience, 28(2), 222–233. 10.1038/s41593-024-01832-y

Ekhtiari, H., Kuplicki, R., Yeh, H., & Paulus, M. P. (2019). Physical characteristics not psychological state or trait characteristics predict motion during resting state fMRI. Scientific Reports, 9(1), 419. 10.1038/s41598-018-36699-0

Esteban, O., Birman, D., Schaer, M., Koyejo, O. O., Poldrack, R. A., & Gorgolewski, K. J. (2017). MRIQC: Advancing the automatic prediction of image quality in MRI from unseen sites. PloS One, 12(9), e0184661. 10.1371/journal.pone.0184661

Fair, D. A., Miranda-Dominguez, O., Snyder, A. Z., Perrone, A., Earl, E. A., Van, A. N., Koller, J. M., Feczko, E., Tisdall, M. D., Van Der Kouwe, A., Klein, R. L., Mirro, A. E., Hampton, J. M., Adeyemo, B., Laumann, T. O., Gratton, C., Greene, D. J., Schlaggar, B. L., Hagler, D. J., … Dosenbach, N. U. F. (2020). Correction of respiratory artifacts in MRI head motion estimates. NeuroImage, 208, 116400. 10.1016/j.neuroimage.2019.116400

Falk, E. B., Hyde, L. W., Mitchell, C., Faul, J., Gonzalez, R., Heitzeg, M. M., Keating, D. P., Langa, K. M., Martz, M. E., Maslowsky, J., et al. (2013). What is a representative brain? Neuroscience meets population science. Proceedings of the National Academy of Sciences, 110(44), 17615–17622. 10.1073/pnas.1310134110

Faraji-Dana, Z., Tam, F., Chen, J. J., & Graham, S. J. (2016a). A robust method for suppressing motion-induced coil sensitivity variations during prospective correction of head motion in fMRI. Magnetic Resonance Imaging, 34(8), 1206–1219. 10.1016/j.mri.2016.06.005

Faraji-Dana, Z., Tam, F., Chen, J. J., & Graham, S. J. (2016b). Interactions between head motion and coil sensitivity in accelerated fMRI. Journal of Neuroscience Methods, 270, 46–60. 10.1016/j.jneumeth.2016.06.005

Feczko, E., Conan, G., Marek, S., Tervo-Clemmens, B., Cordova, M., Doyle, O., Earl, E., Perrone, A., Sturgeon, D., Klein, R., Harman, G., Kilamovich, D., Hermosillo, R., Miranda-Dominguez, O., Adebimpe, A., Bertolero, M., Cieslak, M., Covitz, S., Hendrickson, T., … Fair, D. A. (2021). Adolescent brain cognitive development (ABCD) community MRI collection and utilities. bioRxiv. 10.1101/2021.07.09.451638

Feinberg, D. A., Moeller, S., Smith, S. M., Auerbach, E., Ramanna, S., Glasser, M. F., Miller, K. L., Ugurbil, K., & Yacoub, E. (2010). Multiplexed echo planar imaging for sub-second whole brain FMRI and fast diffusion imaging. PloS One, 5(12), e15710. 10.1371/journal.pone.0015710

Frew, S., Samara, A., Shearer, H., Eilbott, J., & Vanderwal, T. (2022). Getting the nod: Pediatric head motion in a transdiagnostic sample during movie- and resting-state fMRI. PLOS ONE, 17(4), e0265112. 10.1371/journal.pone.0265112

Friston, K. J., Williams, S., Howard, R., & Frackowiak, R. S. J. (1996). Movement-related effects in fMRI time-series. Magnetic Resonance Imaging, 35(3), 346–355. 10.1002/mrm.1910350312

Fry, A., Littlejohns, T. J., Sudlow, C., Doherty, N., Adamska, L., Sprosen, T., Collins, R., & Allen, N. E. (2017). Comparison of Sociodemographic and Health-Related Characteristics of UK Biobank Participants With Those of the General Population. American Journal of Epidemiology, 186(9), 1026–1034. 10.1093/aje/kwx246

Gamer, M., Lemon, J., & Singh, I. F. P. (2019). Irr: Various coefficients of interrater reliability and agreement. https://www.r-project.org

Gao, P., Wang, Y.-S., Lu, Q.-Y., Rong, M.-J., Fan, X.-R., Holmes, A. J., Dong, H.-M., Li, H.-F., & Zuo, X.-N. (2023). Brief mock-scan training reduces head motion during real scanning for children: A growth curve study. Developmental Cognitive Neuroscience, 61, 101244. 10.1016/j.dcn.2023.101244

Garavan, H., Bartsch, H., Conway, K., Decastro, A., Goldstein, R. Z., Heeringa, S., Jernigan, T., Potter, A., Thompson, W., & Zahs, D. (2018). Recruiting the ABCD sample: Design considerations and procedures. Developmental Cognitive Neuroscience, 32, 16–22. 10.1016/j.dcn.2018.04.004

Gard, A. M., Hyde, L. W., Heeringa, S. G., West, B. T., & Mitchell, C. (2023). Why weight? Analytic approaches for large-scale population neuroscience data. Developmental Cognitive Neuroscience, 59, 101196. 10.1016/j.dcn.2023.101196

Glasser, M. F., Coalson, T. S., Bijsterbosch, J. D., Harrison, S. J., Harms, M. P., Anticevic, A., Van Essen, D. C., & Smith, S. M. (2018). Using temporal ICA to selectively remove global noise while preserving global signal in functional MRI data. Neuroimage, 181, 692–717. 10.1016/j.neuroimage.2018.04.076

Greene, A. S., Gao, S., Scheinost, D., & Constable, R. T. (2018). Task-induced brain state manipulation improves prediction of individual traits. Nature Communications, 9(1), 2807. 10.1038/s41467-018-04920-3

Griffanti, L., Salimi-Khorshidi, G., Beckmann, C. F., Auerbach, E. J., Douaud, G., Sexton, C. E., Zsoldos, E., Ebmeier, K. P., Filippini, N., Mackay, C. E., et al. (2014). ICA-based artefact removal and accelerated fMRI acquisition for improved resting state network imaging. Neuroimage, 95, 232–247. 10.1016/j.neuroimage.2014.03.034

Hagler Jr, D. J., Hatton, S., Cornejo, M. D., Makowski, C., Fair, D. A., Dick, A. S., Sutherland, M. T., Casey, B., Barch, D. M., Harms, M. P., et al. (2019). Image processing and analysis methods for the adolescent brain cognitive development study. Neuroimage, 202, 116091. 10.1016/j.neuroimage.2019.116091

Hariri, A. R., Tessitore, A., Mattay, V. S., Fera, F., & Weinberger, D. R. (2002). The Amygdala Response to Emotional Stimuli: A Comparison of Faces and Scenes. NeuroImage, 17(1), 317–323. 10.1006/nimg.2002.1179

Harms, M. P., Somerville, L. H., Ances, B. M., Andersson, J., Barch, D. M., Bastiani, M., Bookheimer, S. Y., Brown, T. B., Buckner, R. L., Burgess, G. C., Coalson, T. S., Chappell, M. A., Dapretto, M., Douaud, G., Fischl, B., Glasser, M. F., Greve, D. N., Hodge, C., Jamison, K. W., … Yacoub, E. (2018). Extending the Human Connectome Project across ages: Imaging protocols for the Lifespan Development and Aging projects. NeuroImage, 183, 972–984. 10.1016/j.neuroimage.2018.09.060

Havsteen, I., Ohlhues, A., Madsen, K. H., Nybing, J. D., Christensen, H., & Christensen, A. (2017). Are movement artifacts in magnetic resonance imaging a real problem?—a narrative review. Frontiers in Neurology, 8, 232. 10.3389/fneur.2017.00232

Health Survey for England. (2022).

Hernán, M. A., Hernández-Dìaz, S., & Robins, J. M. (2004). A structural approach to selection bias. Epidemiology, 15(5), 615–625. 10.1097/01.ede.0000135174.63482.43

Jung, H., Amini, M., Hunt, B. J., Murphy, E. I., Sadil, P., Halchenko, Y. O., Petre, B., Miao, Z., Kragel, P. A., Han, X., et al. (2025). Spacetop: A multimodal fMRI dataset unifying naturalistic processes with a rich array of experimental tasks. Scientific Data, 12(1), 1465. 10.1038/s41597-025-05154-x

Kay, B. P., Montez, D. F., Marek, S., Tervo-Clemmens, B., Siegel, J. S., Adeyemo, B., Laumann, T. O., Metoki, A., Chauvin, R. J., Van, A. N., et al. (2025). Motion impact score for detecting spurious brain-behavior associations. Nature Communications, 16(1), 8614. 10.1038/s41467-025-63661-2

Kelleher, E. M., & Irani, A. (2026). What can UK biobank’s 500 000 participants teach us about chronic pain? In British Journal of Anaesthesia. Elsevier. 10.1016/j.bja.2026.02.027

Keyes, K. M., & Westreich, D. (2019). UK Biobank, big data, and the consequences of non-representativeness. The Lancet, 393(10178), 1297. 10.1016/S0140-6736(18)33067-8

Knutson, B., Westdorp, A., Kaiser, E., & Hommer, D. (2000). FMRI Visualization of Brain Activity during a Monetary Incentive Delay Task. NeuroImage, 12(1), 20–27. 10.1006/nimg.2000.0593

Kong, X., Zhen, Z., Li, X., Lu, H., Wang, R., Liu, L., He, Y., Zang, Y., & Liu, J. (2014). Individual Differences in Impulsivity Predict Head Motion during Magnetic Resonance Imaging. PLoS ONE, 9(8), e104989. 10.1371/journal.pone.0104989

LeWinn, K. Z., Sheridan, M. A., Keyes, K. M., Hamilton, A., & McLaughlin, K. A. (2017). Sample composition alters associations between age and brain structure. Nature Communications, 8(1), 874. 10.1038/s41467-017-00908-7

Liu, W., Wei, D., Chen, Q., Yang, W., Meng, J., Wu, G., Bi, T., Zhang, Q., Zuo, X.-N., & Qiu, J. (2017). Longitudinal test-retest neuroimaging data from healthy young adults in southwest china. Scientific Data, 4(1), 1–9. 10.1038/sdata.2017.17

Logan, G. D. (1994). Spatial attention and the apprehension of spatial relations. Journal of Experimental Psychology: Human Perception and Performance, 20(5), 1015–1036. 10.1037/0096-1523.20.5.1015

Louis, T. A., & Zeger, S. L. (2008). Effective communication of standard errors and confidence intervals. Biostatistics, 10(1), 1–2. 10.1093/biostatistics/kxn014

Maknojia, S., Churchill, N. W., Schweizer, T. A., & Graham, S. J. (2019). Resting State fMRI: Going Through the Motions. Frontiers in Neuroscience, 13, 825. 10.3389/fnins.2019.00825

Makowski, C., Lepage, M., & Evans, A. C. (2019). Head motion: The dirty little secret of neuroimaging in psychiatry. Journal of Psychiatry and Neuroscience, 44(1), 62–68. 10.1503/jpn.180022

Mansour, S. L., Di Biase, M. A., Smith, R. E., Zalesky, A., & Seguin, C. (2023). Connectomes for 40,000 UK biobank participants: A multi-modal, multi-scale brain network resource. NeuroImage, 283, 120407. 10.1016/j.neuroimage.2023.120407

Marcus, D. S., Harms, M. P., Snyder, A. Z., Jenkinson, M., Wilson, J. A., Glasser, M. F., Barch, D. M., Archie, K. A., Burgess, G. C., Ramaratnam, M., Hodge, M., Horton, W., Herrick, R., Olsen, T., McKay, M., House, M., Hileman, M., Reid, E., Harwell, J., … Van Essen, D. C. (2013). Human Connectome Project informatics: Quality control, database services, and data visualization. NeuroImage, 80, 202–219. 10.1016/j.neuroimage.2013.05.077

Marek, S., Tervo-Clemmens, B., Calabro, F. J., Montez, D. F., Kay, B. P., Hatoum, A. S., Donohue, M. R., Foran, W., Miller, R. L., Hendrickson, T. J., et al. (2022). Reproducible brain-wide association studies require thousands of individuals. Nature, 603(7902), 654–660. 10.1038/s41586-022-04492-9

Marek, S., Tervo-Clemmens, B., Nielsen, A. N., Wheelock, M. D., Miller, R. L., Laumann, T. O., Earl, E., Foran, W. W., Cordova, M., Doyle, O., Perrone, A., Miranda-Dominguez, O., Feczko, E., Sturgeon, D., Graham, A., Hermosillo, R., Snider, K., Galassi, A., Nagel, B. J., … Dosenbach, N. U. F. (2019). Identifying reproducible individual differences in childhood functional brain networks: An ABCD study. Developmental Cognitive Neuroscience, 40, 100706. 10.1016/j.dcn.2019.100706

McGillycuddy, M., Popovic, G., Bolker, B. M., & Warton, D. I. (2025). Parsimoniously fitting large multivariate random effects in glmmTMB. Journal of Statistical Software, 112, 1–19. 10.18637/jss.v112.i01

Meissner, T. W., Walbrin, J., Nordt, M., Koldewyn, K., & Weigelt, S. (2020). Head motion during fMRI tasks is reduced in children and adults if participants take breaks. Developmental Cognitive Neuroscience, 44, 100803. 10.1016/j.dcn.2020.100803

Mejia, A., Hwang, J., Pham, D., Noble, S., Satterthwaite, T. D., Nichols, T. E., & Yeo, B. (2026). Excessive data censoring in fMRI undermines individual precision and weakens brain-behavior associations. arXiv Preprint arXiv:2603.07380. 10.48550/arXiv.2603.07380

Miller, K. L., Alfaro-Almagro, F., Bangerter, N. K., Thomas, D. L., Yacoub, E., Xu, J., Bartsch, A. J., Jbabdi, S., Sotiropoulos, S. N., Andersson, J. L. R., Griffanti, L., Douaud, G., Okell, T. W., Weale, P., Dragonu, I., Garratt, S., Hudson, S., Collins, R., Jenkinson, M., … Smith, S. M. (2016). Multimodal population brain imaging in the UK Biobank prospective epidemiological study. Nature Neuroscience, 19(11), 1523–1536. 10.1038/nn.4393

Moeller, S., Yacoub, E., Olman, C. A., Auerbach, E., Strupp, J., Harel, N., & Uğurbil, K. (2010). Multiband multislice GE-EPI at 7 tesla, with 16-fold acceleration using partial parallel imaging with application to high spatial and temporal whole-brain fMRI. Magnetic Resonance in Medicine, 63(5), 1144–1153. 10.1002/mrm.22361

Muschelli, J., Beth, M., Caffo, B. S., Barber, A. D., Pekar, J. J., & Mostofsky, S. H. (2014). NeuroImage reduction of motion-related artifacts in resting state fMRI using aCompCor. NeuroImage, 96, 22–35. 10.1016/j.neuroimage.2014.03.028

Nebel, M. B., Lidstone, D. E., Wang, L., Benkeser, D., Mostofsky, S. H., & Risk, B. B. (2022). Accounting for motion in resting-state fMRI: What part of the spectrum are we characterizing in autism spectrum disorder? NeuroImage, 257, 119296. 10.1016/j.neuroimage.2022.119296

Nichols, T. E., Das, S., Eickhoff, S. B., Evans, A. C., Glatard, T., Hanke, M., Kriegeskorte, N., Milham, M. P., Poldrack, R. A., Poline, J.-B., Proal, E., Thirion, B., Van Essen, D. C., White, T., & Yeo, B. T. T. (2016). Best practices in data analysis and sharing in neuroimaging using MRI. bioRxiv. 10.1101/054262

Ooi, L. Q. R., Orban, C., Zhang, S., Nichols, T. E., Tan, T. W. K., Kong, R., Marek, S., Dosenbach, N. U., Laumann, T. O., Gordon, E. M., et al. (2025). Longer scans boost prediction and cut costs in brain-wide association studies. Nature, 1–10. 10.1038/s41586-025-09250-1

Pardoe, H. R., Hiess, R. K., & Kuzniecky, R. (2016). Motion and morphometry in clinical and nonclinical populations. Neuroimage, 135, 177–185. 10.1016/j.neuroimage.2016.05.005

Parkes, L., Fulcher, B., Yücel, M., & Fornito, A. (2018). An evaluation of the efficacy, reliability, and sensitivity of motion correction strategies for resting-state functional MRI. NeuroImage, 171, 415–436. 10.1016/j.neuroimage.2017.12.073

Peverill, M., Russell, J. D., Keding, T. J., Rich, H. M., Halvorson, M. A., King, K. M., Birn, R. M., & Herringa, R. J. (2025). Balancing data quality and bias: Investigating functional connectivity exclusions in the adolescent brain cognitive development℠(ABCD study) across quality control pathways. Human Brain Mapping, 46(1), e70094. 10.1002/hbm.70094

Phạm, D., McDonald, D. J., Ding, L., Nebel, M. B., & Mejia, A. F. (2023). Less is more: Balancing noise reduction and data retention in fMRI with data-driven scrubbing. NeuroImage, 270, 119972.

Power, J. D., Barnes, K. A., Snyder, A. Z., Schlaggar, B. L., & Petersen, S. E. (2012). Spurious but systematic correlations in functional connectivity MRI networks arise from subject motion. NeuroImage, 59(3), 2142–2154. 10.1016/j.neuroimage.2011.10.018

Power, J. D., Lynch, C. J., Adeyemo, B., & Petersen, S. E. (2020). A critical, event-related appraisal of denoising in resting-state fMRI studies. Cerebral Cortex, 30(10), 5544–5559. 10.1093/cercor/bhaa139

Power, J. D., Lynch, C. J., Silver, B. M., Dubin, M. J., Martin, A., & Jones, R. M. (2019). Distinctions among real and apparent respiratory motions in human fMRI data. NeuroImage, 201, 116041. 10.1016/j.neuroimage.2019.116041

Power, J. D., Mitra, A., Laumann, T. O., Snyder, A. Z., Schlaggar, B. L., & Petersen, S. E. (2014). Methods to detect, characterize, and remove motion artifact in resting state fMRI. NeuroImage, 84(Supplement C), 320–341. 10.1016/j.neuroimage.2013.08.048

Pruim, R. H. R., Mennes, M., Van Rooij, D., Llera, A., Buitelaar, J. K., & Beckmann, C. F. (2015). ICA-AROMA: A robust ICA-based strategy for removing motion artifacts from fMRI data. NeuroImage, 112, 267–277. 10.1016/j.neuroimage.2015.02.064

Ramduny, J., Uddin, L. Q., Vanderwal, T., Feczko, E., Fair, D. A., Kelly, C., & Baskin-Sommers, A. (2025). Representing brain-behavior associations by retaining high-motion minoritized youth. Biological Psychiatry: Cognitive Neuroscience and Neuroimaging. 10.1016/j.bpsc.2025.01.014

Reddy, N. A., Zvolanek, K. M., Moia, S., Caballero-Gaudes, C., & Bright, M. G. (2023). Denoising task-correlated head motion from motor-task fMRI data with multi-echo ICA. bioRxiv: The Preprint Server for Biology, 2023.07.19.549746. 10.1101/2023.07.19.549746

Ricard, J., Parker, T., Dhamala, E., Kwasa, J., Allsop, A., & Holmes, A. (2023). Confronting racially exclusionary practices in the acquisition and analyses of neuroimaging data. Nature Neuroscience, 26(1), 4–11. 10.1038/s41593-022-01218-y

Satterthwaite, T. D., Wolf, D. H., Ruparel, K., Erus, G., Elliott, M. A., Eickhoff, S. B., Gennatas, E. D., Jackson, C., Prabhakaran, K., Smith, A., Hakonarson, H., Verma, R., Davatzikos, C., Gur, R. E., & Gur, R. C. (2013). Heterogeneous impact of motion on fundamental patterns of developmental changes in functional connectivity during youth. NeuroImage, 83, 45–57. 10.1016/j.neuroimage.2013.06.045

Schaefer, A., Kong, R., Gordon, E. M., Laumann, T. O., Zuo, X.-N., Holmes, A. J., Eickhoff, S. B., & Yeo, B. T. (2018). Local-global parcellation of the human cerebral cortex from intrinsic functional connectivity MRI. Cerebral Cortex, 28(9), 3095–3114. 10.1093/cercor/bhx179

Seto, E., Sela, G., McIlroy, W. E., Black, S. E., Staines, W. R., Bronskill, M. J., McIntosh, A. R., & Graham, S. J. (2001). Quantifying Head Motion Associated with Motor Tasks Used in fMRI. NeuroImage, 14(2), 284–297. 10.1006/nimg.2001.0829

Setsompop, K., Gagoski, B. A., Polimeni, J. R., Witzel, T., Wedeen, V. J., & Wald, L. L. (2012). Blipped-controlled aliasing in parallel imaging for simultaneous multislice echo planar imaging with reduced g-factor penalty. Magnetic Resonance in Medicine, 67(5), 1210–1224. 10.1002/mrm.23097

Shi, T. C., Durham, K., Marsh, R., & Pagliaccio, D. (2025). Differences in head motion during functional magnetic resonance imaging across pediatric neuropsychiatric disorders. Biological Psychiatry Global Open Science, 5(3), 100446. 10.1016/j.bpsgos.2024.100446

Siegel, J. S., Mitra, A., Laumann, T. O., Seitzman, B. A., Raichle, M., Corbetta, M., & Snyder, A. Z. (2017). Data Quality Influences Observed Links Between Functional Connectivity and Behavior. Cerebral Cortex, 27(9), 4492–4502. 10.1093/cercor/bhw253

Somerville, L. H., Bookheimer, S. Y., Buckner, R. L., Burgess, G. C., Curtiss, S. W., Dapretto, M., Elam, J. S., Gaffrey, M. S., Harms, M. P., Hodge, C., et al. (2018). The lifespan human connectome project in development: A large-scale study of brain connectivity development in 5–21 year olds. Neuroimage, 183, 456–468. 10.1016/j.neuroimage.2018.08.050

Teo, T. W. J., Saffari, S. E., Chan, L. L., & Welton, T. (2024). Comparison of MRI head motion indicators in 40,969 subjects informs neuroimaging study design. Scientific Reports, 14(1), 29430. 10.1038/s41598-024-79827-9

Thompson, W. K., Barch, D. M., Bjork, J. M., Gonzalez, R., Nagel, B. J., Nixon, S. J., & Luciana, M. (2019). The structure of cognition in 9 and 10 year-old children and associations with problem behaviors: Findings from the ABCD study’s baseline neurocognitive battery. Developmental Cognitive Neuroscience, 36, 100606. 10.1111/nyas.14268

Van De Moortele, P.-F., Pfeuffer, J., Glover, G. H., Ugurbil, K., & Hu, X. (2002). Respiration-induced *B*_0_ fluctuations and their spatial distribution in the human brain at 7 Tesla. Magnetic Resonance in Medicine, 47(5), 888–895. 10.1002/mrm.10145

Van Dijk, K. R., Sabuncu, M. R., & Buckner, R. L. (2012). The influence of head motion on intrinsic functional connectivity MRI. Neuroimage, 59(1), 431–438. 10.1016/j.neuroimage.2011.07.044

Ward, J., Cox, S. R., Quinn, T., Lyall, L. M., Strawbridge, R. J., Russell, E., Pell, J. P., Stewart, W., Cullen, B., Whalley, H., & Lyall, D. M. (2024). Head motion in the UK Biobank imaging subsample: Longitudinal stability, associations with psychological and physical health, and risk of incomplete data. Brain Communications, 6(4), fcae220. 10.1093/braincomms/fcae220

Weinberger, D. R., & Radulescu, E. (2016). Finding the elusive psychiatric “lesion” with 21st-century neuroanatomy: A note of caution. American Journal of Psychiatry, 173(1), 27–33. 10.1176/appi.ajp.2015.15060753

White, N., Roddey, C., Shankaranarayanan, A., Han, E., Rettmann, D., Santos, J., Kuperman, J., & Dale, A. (2010). PROMO: Real-time prospective motion correction in MRI using image-based tracking. Magnetic Resonance in Medicine: An Official Journal of the International Society for Magnetic Resonance in Medicine, 63(1), 91–105. 10.1002/mrm.22176

Williams, J. C., Tubiolo, P. N., Luceno, J. R., & Van Snellenberg, J. X. (2022). Advancing motion denoising of multiband resting-state functional connectivity fMRI data. Neuroimage, 249, 118907. 10.1016/j.neuroimage.2022.118907

Wylie, G. R., Genova, H., DeLuca, J., Chiaravalloti, N., & Sumowski, J. F. (2014). Functional magnetic resonance imaging movers and shakers: Does subject-movement cause sampling bias?: Subject Motion and Sampling Bias. Human Brain Mapping, 35(1), 1–13. 10.1002/hbm.22150

Xu, J., Moeller, S., Strupp, J., Auerbach, E., Chen, L., Feinberg, D. A., Ugurbil, K., & Yacoub, E. (2012). Highly accelerated whole brain imaging using aligned-blipped-controlled-aliasing multiband EPI. Proceedings of the 20th Annual Meeting of ISMRM, 2306, 1907–1913.

Yan, C.-G., Cheung, B., Kelly, C., Colcombe, S., Craddock, R. C., Di Martino, A., Li, Q., Zuo, X.-N., Castellanos, F. X., & Milham, M. P. (2013). A comprehensive assessment of regional variation in the impact of head micromovements on functional connectomics. NeuroImage, 76, 183–201. 10.1016/j.neuroimage.2013.03.004

Yang, C., Coalson, T. S., Smith, S. M., Elam, J. S., Van Essen, D. C., & Glasser, M. F. (2024). Automating the Human Connectome Project’s Temporal ICA Pipeline. 10.1101/2024.01.15.574667

Yao, N., Winkler, A. M., Barrett, J., Book, G. A., Beetham, T., Horseman, R., Leach, O., Hodgson, K., Knowles, E. E., Mathias, S., et al. (2017). Inferring pathobiology from structural MRI in schizophrenia and bipolar disorder: Modeling head motion and neuroanatomical specificity. Human Brain Mapping, 38(8), 3757–3770. 10.1002/hbm.23612

Zeng, L.-L., Wang, D., Fox, M. D., Sabuncu, M., Hu, D., Ge, M., Buckner, R. L., & Liu, H. (2014). Neurobiological basis of head motion in brain imaging. Proceedings of the National Academy of Sciences, 111(16), 6058–6062. 10.1073/pnas.1317424111

Zuo, X.-N., Anderson, J. S., Bellec, P., Birn, R. M., Biswal, B. B., Blautzik, J., Breitner, J., Buckner, R. L., Calhoun, V. D., Castellanos, F. X., et al. (2014). An open science resource for establishing reliability and reproducibility in functional connectomics. Scientific Data, 1(1), 1–13. 10.1038/sdata.2014.49

